# Neural circuit mechanisms for multi-tasking

**DOI:** 10.64898/2026.10.01.755960

**Authors:** Manuel Beiran, Owen Marschall, Klavdia Zemlianova, Arianna Di Bernardo, Srdjan Ostojic, Ashok Litwin-Kumar

## Abstract

Neural activity during performance of a cognitive or motor task is often described as lying on a low-dimensional manifold in the space of all possible activity patterns. When multiple tasks are performed, an individual brain region’s activity may generate multiple such manifolds. Several models have been introduced to explain this capability, each with different assumptions concerning architecture and inputs. Here, we develop a theory of multi-tasking according to which models and experimental paradigms can be organized. We introduce a distinction between sequential and simultaneous multi-tasking, the performance of one task at a time or multiple in parallel. We show that interference between tasks often prevents multi-tasking and introduce a model of “task barcodes” that circumvents this problem and parameterizes a neuronal population’s task selectivity. Our theory reveals categorical distinctions between neural circuit mechanisms supporting multi-tasking that can be probed using physiological recordings.

## Introduction

The ability to generate different responses across changing goals and circumstances is a defining feature of intelligent behavior. This ability requires neural circuits to flexibly map sensory inputs to actions, across conditions with distinct behavioral demands. How such multi-tasking is achieved, without interference across conditions, remains an open question.

Simultaneous recordings of many neurons in animals trained to perform multiple tasks have characterized the geometry of neural activity associated with this behavioral flexibility. In primates ^1–4^ and rodents^5,6^, different tasks, defined by specific combinations of stimuli and responses, are often associated with neural activity that lies in distinct regions of neural firing rate space. These regions are sometimes termed “task manifolds” ^7,8^. Here, we ask how a single network can multi-task, that is, support multiple task manifolds each with different associated dynamics.

Numerous modeling studies have addressed aspects of this question, using different assumptions on the neural architecture that supports multi-tasking. Recurrent neural networks trained on multiple tasks frequently rely on explicit contextual inputs that modify a network’s dynamics depending on task demands^9–11^. Other studies have emphasized the role of preparatory activity that instructs a network to produce a specific behavior by setting an appropriate initial condition ^12,13^. Theoretical studies have examined how changes in effective connectivity, mediated by mechanisms such as gating or synaptic modulation, can change a network’s dynamics^14–16^. Related ideas have also been explored in models of continual learning^17^ and contextual inference^18^. What are the relative advantages and limitations of each of these proposed mechanisms, and is each suitable for only certain behaviors or for any behavior that requires multi-tasking?

Here, we develop a theory that characterizes strategies a single recurrent network may employ to achieve multi-tasking. We propose a formal definition of what it means for a single network to multi-task without interference, and show that such interference frequently occurs for nonlinear recurrent neural networks. Our theory introduces a distinction between sequential multi-tasking, in which one task is performed at a time, and simultaneous multi-tasking, in which multiple are performed in parallel. We demonstrate that previous models of multi-tasking can be organized based on their mechanism for switching between tasks and avoiding interference. We also introduce a new model of “task barcodes” that limits interference and enables both simultaneous and sequential multi-tasking. Our analysis of this model identifies a novel relationship between the distribution of single-neuron task selectivities, which can be measured with neural population recordings, and a network’s ability to multi-task. We analyze concrete implementations of multi-tasking mechanisms and show that each produces qualitatively different features in neural firing rate dynamics. Finally, we describe how these mechanisms may be distinguished using neural data.

## Results

### Task variables and types of multi-tasking

Numerous studies have formalized the neural computations that support executing a single sensorimotor task, or solving a cognitive task, as dynamics operating on a set of *task variables* ^9,10,19–21^ (Fig. 1A). These variables refer to quantities that are represented by neural activity, may evolve over time, and are required for task performance. While some variables may correspond to features of sensory inputs, others may be internally generated, such as neural representations of accumulated evidence, elapsed time, or goals. The ensemble of activity patterns that a neural system generates during one task is often described as lying on a low-dimensional *task manifold* in the space of neural firing rates, with a location on the manifold parameterized by the task variables (Fig. 1A, right). The dynamics of neural activity while the task is performed are represented by the flow field on this manifold. This geometrical language for describing the neural basis of task-related computation has become commonplace across theoretical and systems neuroscience^7,19,22,23^.

**Figure 1.**
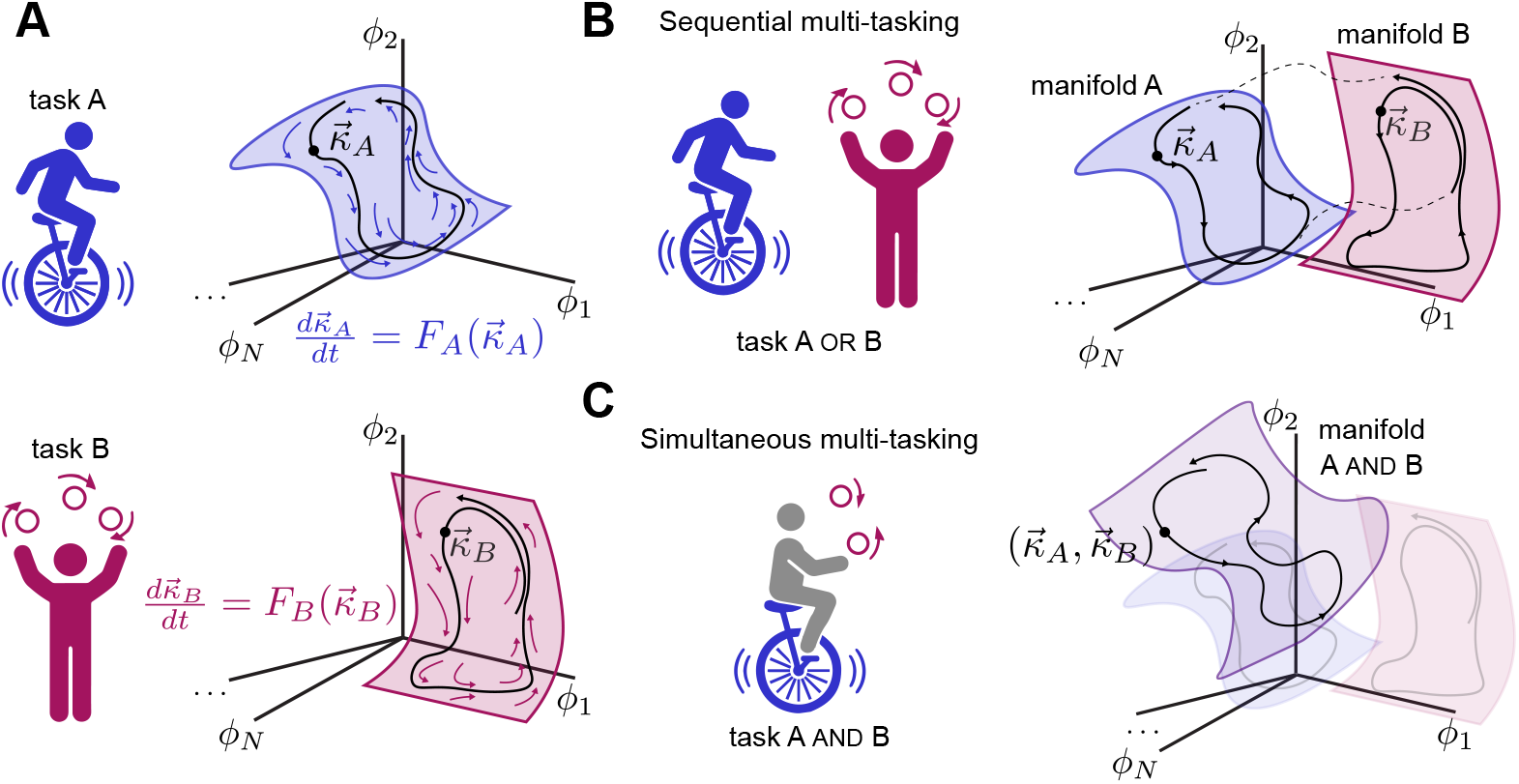
Task manifolds and multi-tasking. **A** Tasks, like cycling or juggling (left) are supported by computations that depend on dynamical interactions between task variables, denoted 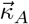 or 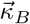 for tasks A and B. These variables may be represented by neural activity that lies on a manifold in firing rate space (right; *ϕ*_*i*_ denotes firing rate of neuron *i*) which is parameterized by 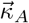 or 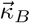. The task variables evolve according to dynamics 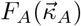 and 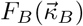 that correspond to flow fields on each manifold (colored arrows). **B** Sequential multi-tasking requires performing one task at a time (left). This may be supported by neural activity that transitions between manifolds associated with each task and is thus parameterized by one set of task variables at a time (right). Parts of trajectories associated with transitions between manifolds are shown with dashed lines. **C** Simultaneous multi-tasking involves performance of multiple tasks at the same time (left). This may be supported by neural activity that is not confined to the manifold corresponding to any one task, but to a new, higher-dimensional manifold that combines them (purple region schematizes a 4-dimensional manifold). For two tasks A and B, the manifold is thus parameterized by 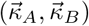.

Individual neural circuits may support multiple sensorimotor tasks, for instance cycling and juggling (Fig. 1A). Each of these requires distinct processing of inputs, such as proprioceptive, vestibular, visual, or somatosensory signals, to generate motor commands. How does the picture of low-dimensional task manifolds extend to networks that multi-task? To answer this, we first distinguish between two types of multi-tasking. One is *sequential* multi-tasking, in which only one task is performed at a time, for example cycling at one moment and juggling at another (Fig. 1B, left). This notion is commonly applied when studying recurrent neural networks trained to perform one of a set of cognitive tasks at a time, given a task-specific contextual cue^9,10^. The second is *simultaneous* multi-tasking, which requires generating outputs for multiple tasks at the same time, for instance cycling and juggling at once (Fig. 1C, left). While this case is under-explored in theoretical studies, complex tasks often require simultaneously solving component subtasks, and some experimental paradigms explicitly demand simultaneous multi-tasking ^24^.

In neural activity space, sequential multi-tasking corresponds to trajectories that switch from one manifold to another (Fig. 1B, right). Thus, the sequential multi-tasking manifold consists of a union of single-task manifolds with their associated flow fields. In contrast, for simultaneous multi-tasking, neural activity is not restricted to any single-task manifold. Instead, it represents all variables across the tasks currently being performed. For example, if tasks A and B are each defined by two variables, activity during the simultaneous performance of A and B lies on a four-dimensional manifold (Fig. 1C, right). Thus, the topology of the simultaneous multi-tasking manifold is that of the Cartesian product of the single-task manifolds. As we will show, different neural architectures may generate manifolds that support these two forms of multi-tasking.

### Implementing tasks through recurrent connectivity

Networks of interconnected neurons may implement task-related computations by generating dynamics through their recurrent interactions^19^. Recurrent neural networks are a popular model class for testing this hypothesis^25,26^, as they can, in principle, approximate arbitrary dynamical systems^27^. In these models, *N* neurons indexed by *i* = 1 … *N* are described by voltage-like variables *x*_*i*_ and communicate through firing rates *ϕ*(*x*_*i*_), where *ϕ*(*x*) is the activation function that maps *x* to firing rate (for our simulations, taken to be a rectified-linear function). The dynamics of individual neurons obey:

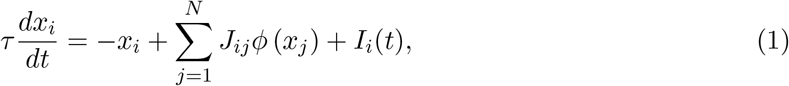

where *J*_*ij*_ is the strength of the connection from neuron *j* to neuron *i, I*_*i*_ is the external input received by neuron *i*, and *τ* is the membrane time constant.

For a network that performs a single task, a particularly direct relation between recurrent connectivity and low-dimensional task dynamics is obtained if the connectivity is assumed to take the form of a low-rank matrix^20,21,28^:

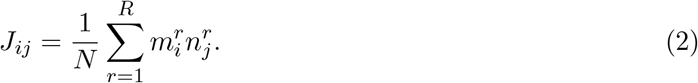

In this case, the dynamics defined by Eq. (1) lie in a subspace whose dimension is at most *R* + *R*_input_ (where *R* is the rank of the matrix and *R*_input_ is the dimension of the external input), which we call the *task subspace* (Fig. 2A, top). Activity is always attracted to this subspace if a network performs only one task. Transforming this activity through the firing rate nonlinearity produces a nonlinear task manifold whose intrinsic dimension equals that of the task subspace (Fig. 2A, bottom). Neural activity within the task subspace, or manifold, represents the variables 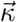 and is the substrate for task-related computations (Fig. 1A). Linear readouts of this activity can produce time-varying outputs that represent solutions to cognitive or motor tasks^20^. Low-rank network models therefore provide an analytically tractable account of neural computation on low-dimensional subspaces (when measuring voltage) or manifolds (when measuring firing rate).

**Figure 2.**
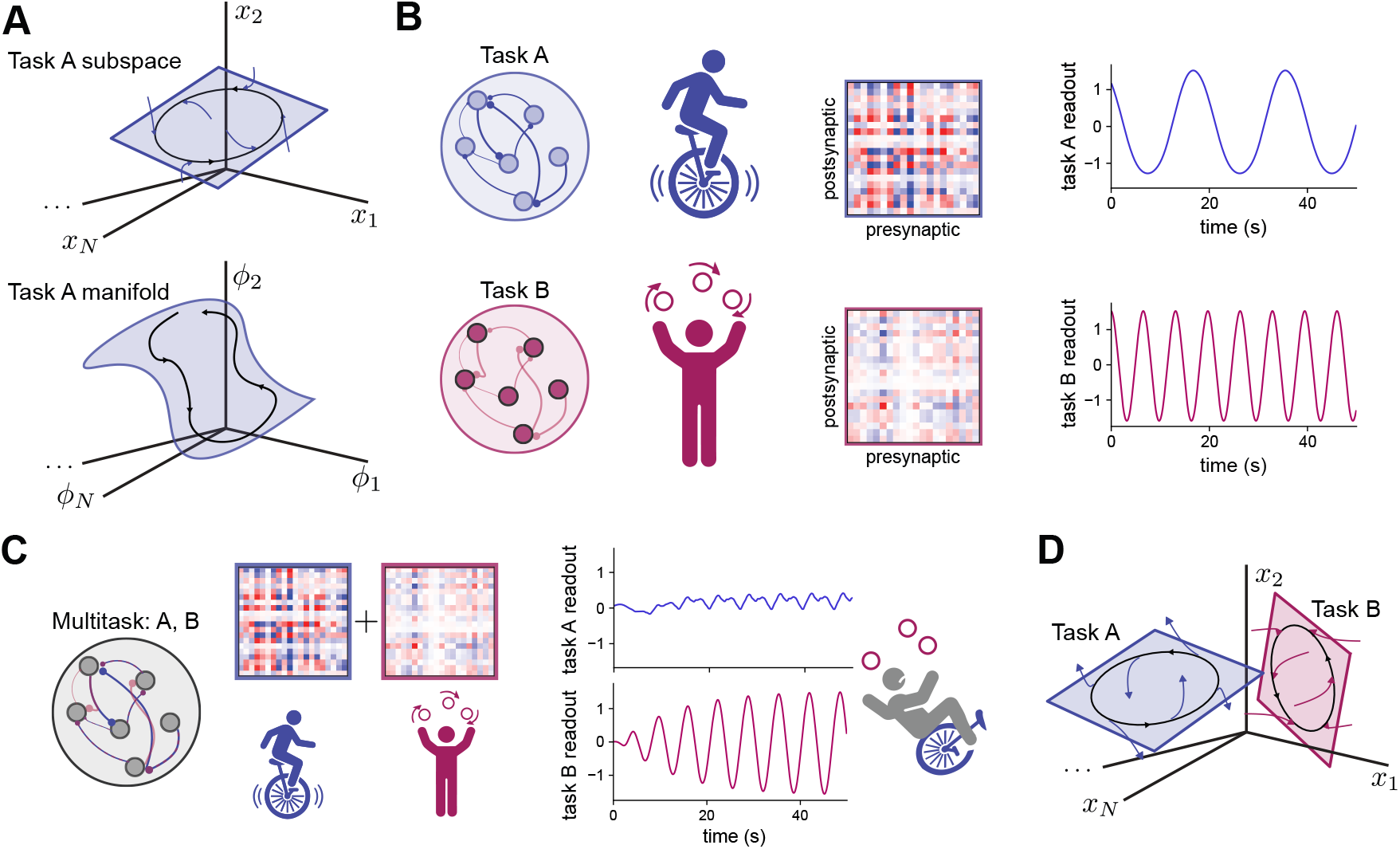
Task interference. **A** Illustration of a two-dimensional task subspace in the space of input currents *x*_*i*_, *i* = 1 … *N* (top) and a two-dimensional task manifold in the space of firing rates *ϕ*_*i*_ (bottom). Blue arrows in top panel indicate the subspace’s transverse stability: trajectories that are perturbed off the subspace are attracted to it. **B** Two networks with rank-two synaptic weights each produce oscillations with different frequencies (right). These oscillations model the computation each network performs in each of two different tasks, in this case cycling (top) or juggling (bottom). **C** A rank-four network whose synaptic weights are the sum of those in **B** (left) fails to multi-task: the task A output is suppressed by task B and occurs at an incorrect frequency (right). **D** Illustration of transverse instability of the task A subspace. Trajectories that are perturbed off the subspace are repelled from it and attracted to the task B subspace.

### Multi-tasking in recurrent neural networks and the problem of interference

To construct a multi-tasking network, we assume that ***J*** is a weighted sum of connectivity components related to different tasks and a non-task-related term that represents global inhibition (see Methods):

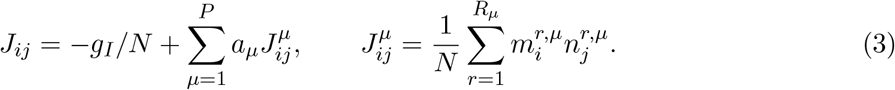

Here, *P* is the number of tasks, *a*_*µ*_ is the strength of the connectivity component associated with task *μ*, and *g*_*I*_ is the magnitude of global inhibition. Each component ***J*** ^*µ*^ is a rank-*R*_*µ*_ synaptic weight matrix, described by the connectivity vectors ***m***^*r,µ*^ and ***n***^*r,µ*^, representing a task associated with a manifold of dimension *R*_*µ*_ + *R*_input_. The assumption of summed connectivity components underlies the construction of many models of multi-task networks^13,15,29,30^. It can be traced to Hopfield networks, whose weight matrix is a sum of rank-one terms, each corresponding to an individual item to be memorized^31^. A similar structure arises naturally in models of sequential learning, where ***J*** ^*µ*^ is the change in weights produced by learning of the *μ*th task^32,33^.

The ability of a network to multi-task depends on how variables associated with different tasks interact. We begin by considering two immediate examples of recurrent neural networks that lack such interactions completely. The first is a network containing disjoint subsets of neurons, with each subset dedicated to a single task and lacking connections to any other. Such a model is at odds with the common observation of individual neurons exhibiting mixed selectivity to multiple tasks^7,34^. We therefore do not focus on this extreme further (although we will consider a relaxed form of task-specific selectivity later). The second example is a linear network in which each task is associated with an orthogonal subspace of firing rates, generated by orthogonal sets of connectivity vectors across tasks. In this situation, the dynamics within each subspace decouple, factorizing into single-task dynamics much like in networks with disjoint populations (see Appendix). However, the range of possible dynamics in linear networks is limited and does not include, for instance, multistability or asymptotically stable oscillations.

For nonlinear networks, interactions between task variables generically occur, even if the subspaces associated with different tasks are orthogonal (see Appendix). How do these interactions affect the multi-tasking ability of nonlinear networks whose connectivity is defined by Eq. (3)? To examine this, we constructed two-task networks whose connectivity vectors define orthogonal subspaces. Each connectivity component alone produces a limit cycle of a different frequency in the respective subspace (Fig. 2B). For a network whose synaptic weight matrix is the sum of these two components, it may seem that the dynamics in each subspace should decouple, as they do for linear networks. However, such a network produces only one oscillation in the task B subspace, while activity in the task A subspace is suppressed and occurs at an incorrect frequency (Fig. 2C). We use the term *task interference* to refer to interactions between task variables that prevent multi-tasking and say that task B interferes with task A in this example. Providing an external input aligned with either of the task subspaces does not prevent this limitation (Supp. Fig. S1).

For a network whose weight matrix is a sum of task-related components (Eq. 3), interference occurs if the connectivity vectors for each task are non-orthogonal, as expected. More surprisingly however, our mathematical analysis shows that even for orthogonal connectivity vectors, interactions between task variables often cause one or more task subspaces to become unstable. Thus, even if the dynamics constrained to the task A subspace are unchanged, small fluctuations in task B’s variables are amplified and subsequently suppress activity in the task A subspace, as in the example above. We call this type of interference a *transverse instability* of the task A subspace, as it is associated with unstable growth in the orthogonal direction corresponding to the task B subspace (Fig. 2D; compare with 2A, top). In the Appendix, we provide formal criteria for the existence and transverse stability of multi-tasking manifolds, and an analysis of these properties in networks of the form of Eq. (3). Whether a network satisfies these criteria depends on the structure of the weights and the activation function, and the analysis shows that straightforward extensions of nonlinear, low-rank recurrent neural networks to the multi-task setting (e.g. Fig. 2C, and more broadly any models in which the connectivity vectors are jointly Gaussian) generically lead to transverse instability. In subsequent sections, we describe neural circuit mechanisms that circumvent this general problem of task interference.

### Mechanisms for multi-tasking

#### Transient task-related activity

In the network of Fig. 2C, transverse instability implies that steady state activity occupies only one task manifold. One solution to this problem is to concern ourselves not with steady state activity, but instead only with activity that begins at some initial condition and subsequently evolves for a limited time. Selecting a task, then, requires setting an initial condition that transiently produces activity near the desired manifold ^12^.

To more easily visualize the geometry of neural activity associated with multi-tasking, we studied a two-task setting in which task A requires the generation of a limit cycle as before and task B requires bistable dynamics (modeling, for instance, delay period activity following a cue taking one of two values that must be remembered; Fig. 3A). In this case, neural activity can be projected into a three-dimensional space, where the first two axes correspond to the task A subspace, and the third axis corresponds to the task B subspace. However, all of our results generalize to other combinations of tasks, and to different numbers of tasks (see Appendix).

**Figure 3.**
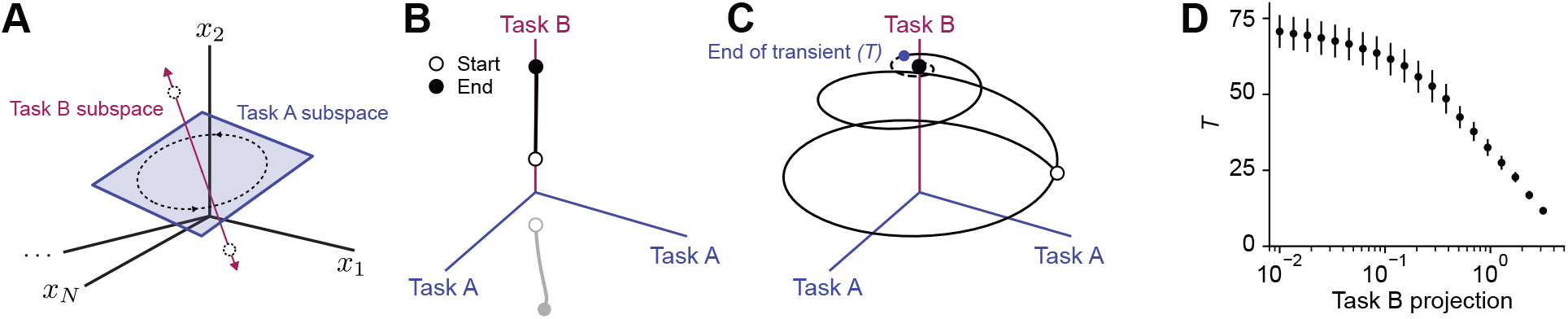
Sequential multi-tasking through transient activity. **A** A two-dimensional task A subspace is associated with a limit cycle (dashed ellipse) and a one-dimensional task B subspace is associated with bistable fixed points (dashed circles). **B** Two neural trajectories (black and gray) projected onto the three-dimensional space defined by the subspaces in **A**. Each initial condition (open circles) lies near the task B subspace and produces a trajectory that approaches one of the two fixed points by the end of the simulation (filled circles). **C** Similar to **B** but for a neural trajectory with initial condition near the task A subspace. The trajectory transiently oscillates before decaying to a fixed point in the task B subspace. Blue mark indicates the time point *T* at which the norm of the projection onto the task A subspace decays below *r* = 0.25 times its initial value. Trajectory after this timepoint is illustrated with a dashed line. **D** Length of transient *T* as defined in **C**, in units of membrane time constant, as a function of the initial condition’s projection onto the task B subspace (relative to that for task A). Mean *±*s.e.m. over 20 networks with *N* = 10,000.

Neural trajectories whose initial conditions overlap with the task B subspace approach one of the two fixed points, as desired (Fig. 3B). However, a limit cycle is produced only if the initial condition is perfectly aligned with the task A subspace. Any small deviation in initial condition along the task B subspace leads to movement toward one of the fixed points and a decaying oscillation. This is due to interference that produces transverse instability of the task A subspace (Fig. 3C; similar to Fig. 2D). We estimated how long task A is performed by measuring the time interval over which the projection of activity on the task A subspace exceeds some fraction of its initial value (Fig. 3C, blue dot). This interval grows as the initial condition is fine-tuned to minimize overlap with the task B subspace (Fig. 3D). However, the growth is at best logarithmic. Thus, extremely precise control of the initial condition is required to perform multiple tasks over long intervals.

Previous models of the production of multiple sequences of neural activity can be viewed as examples of transient multi-tasking, with the length of the recalled sequence proportional to the time interval over which the task is performed. Gillett et al. ^13^ analyzed a model in which the *r*th element in sequence *μ* is associated with an activity pattern ***m***^*µ,r*^ and the connectivity component for each sequence is:

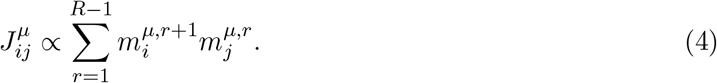

This equation is a special case of Eq. 3, where each sequence *μ* is interpreted as a task and associated with activity in an *R*-dimensional manifold. An initial condition that overlaps with the first element of a sequence triggers the recall of the subsequent elements. However, as above, small overlaps with elements of other sequences produce interference, corresponding to the transverse instability of individual task subspaces^13,35^ (Supp. Fig. S2).

We conclude that transient solutions to sequential multi-tasking are viable for short time intervals, but longer time intervals require increasingly precise control of the initial condition by a preparatory process. Moreover, simultaneous multi-tasking is not easily achieved because an initial condition that has high overlap with multiple task subspaces leads to the task with the greatest degree of amplification quickly suppressing the others. We next investigate mechanisms that reduce this sensitivity to initial state.

#### Multistable task manifolds

Certain network models do not exhibit transverse instability and instead feature multiple stable task manifolds. Such multistable dynamics can be used as the substrate for sequential multi-tasking, with an initial condition set by a preparatory signal shifting the network state from the basin of attraction of one task manifold to another. Classical attractor models of place representations in multiple environments are an example of this^36–38^. In a single environment, place cells provide a spatial map commonly modeled as a continuous attractor. For a one-dimensional environment, the neural representation of space forms a manifold parameterized by one task variable. As an example, we assume a periodic encoding that forms a ring attractor^39^, analogous to the periodic encoding of two-dimensional environments by grid cells ^40^. Place cells can remap across environments, and one hypothesis for this phenomenon is that a map of each environment is represented by an independent continuous attractor (Fig. 4A). A network that generates multiple maps can be obtained by summing weights corresponding to each map^36–38,41^.

**Figure 4.**
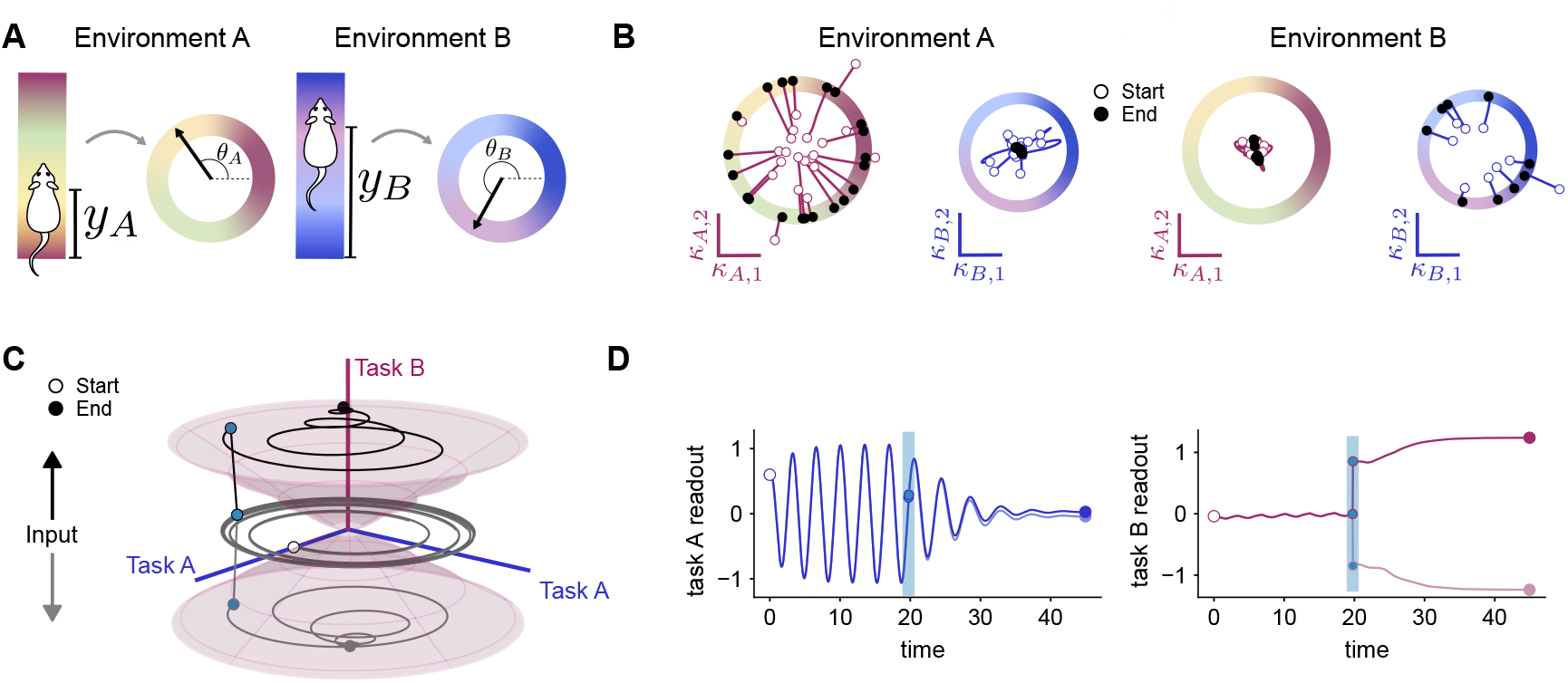
Multistability for two examples of sequential multi-tasking: **A-B** a model of place cells in two one-dimensional environments; **C-D** and a multi-stable combination of a limit cycle and bistable fixed points. **A** An animal represents its location *y*_*A/B*_ in either of two one-dimensional environments (A or B) with a periodic encoding parameterized by *θ*_*A/B*_. **B** Dynamics of neural activity projected onto the subspace associated with each environment. Dynamics in each subspace feature a ring attractor. Locations on the attractor encode *θ*_*A/B*_. Left: Neural trajectories whose initial conditions (open circles) have high overlap with the environment A subspace converge toward a steady state on its corresponding ring (filled circles), while activity in the task B subspace decays. Right: The converse is true for initial conditions that have high overlap with the environment B subspace. **C** A multi-stable model of the limit cycle and memory tasks introduced in Fig. 3. In this model, elements of connectivity vectors follow statistics of a cosine bump, in contrast with Gaussian-distributed weights as in Fig. 3. Two neural trajectories projected onto the three-dimensional space defined as in Fig. 3. The initial condition first selects task A, then an input pulse positively (black) or negatively (gray) aligned with the task B subspace moves the trajectory into task B’s basin of attraction. Conical surfaces denote the separatrix between the task A and B basins of attraction. **D** Readouts of the task A and B subspaces as a function of time, for the trajectories in **C**. Blue vertical line indicates when the input pulse is provided.

This can be viewed as an example of multi-tasking in which each task requires a persistent representation of a location on a different map. In its simplest version, a ring attractor network with cosine-tuned synaptic weights is a rank-two recurrent neural network^42^, and adding weights of two such networks produces a rank-four network^38^. An analysis using the theory of low-rank networks shows that, for a range of parameters, this rank-four network exhibits bistability between two ring attractors embedded in mutually orthogonal subspaces ^29^. Depending on the initial condition, activity converges to one of the two manifolds, an example of sequential multi-tasking (Fig. 4B).

This mechanism to generate multistability can also be applied to dynamic tasks (Fig. 4C). For this example, we studied the limit cycle and memory tasks of Fig. 3. Unlike the previous section, multistability requires a different construction of connectivity vectors: instead of Gaussian statistics, the elements of each vector follow the statistics of a cosine bump, as in ring attractor models ^29^ (see Methods). This produces a network for which each task subspace has a basin of attraction in its vicinity (Fig. 4C). Preparatory inputs can then select tasks by moving the network state from one basin of attraction to another (Fig. 4D).

The cosine-tuned connectivity described above is a specific example of a synaptic weight distribution that prevents task interference, but how to avoid interference outside of this case is not yet clear. We developed a theory of the effect of synaptic weight distribution on the ability to sequentially or simultaneously multi-task that predicts the effects of cosine-tuned connectivity components (see Appendix). The theory also led us to introduce a simple mechanism that can be applied to arbitrary weight distributions to mitigate interference, which we will describe in a later section.

#### Modulation of effective connectivity

Other previous models of sequential multi-tasking have proposed modulation of the synaptic weight matrix as a mechanism for task selection^15,18,30,43^. In these models, it is assumed that task selection involves adjusting the coefficients *a*_*µ*_ in Eq. (3) to increase or decrease the strengths of connectivity components associated with different tasks (Fig. 5A). Unlike the previous sections, this mechanism implies that the flow field describing the evolution of neural firing rates changes upon the selection of a new task, so that selecting a task stabilizes the corresponding manifold.

**Figure 5.**
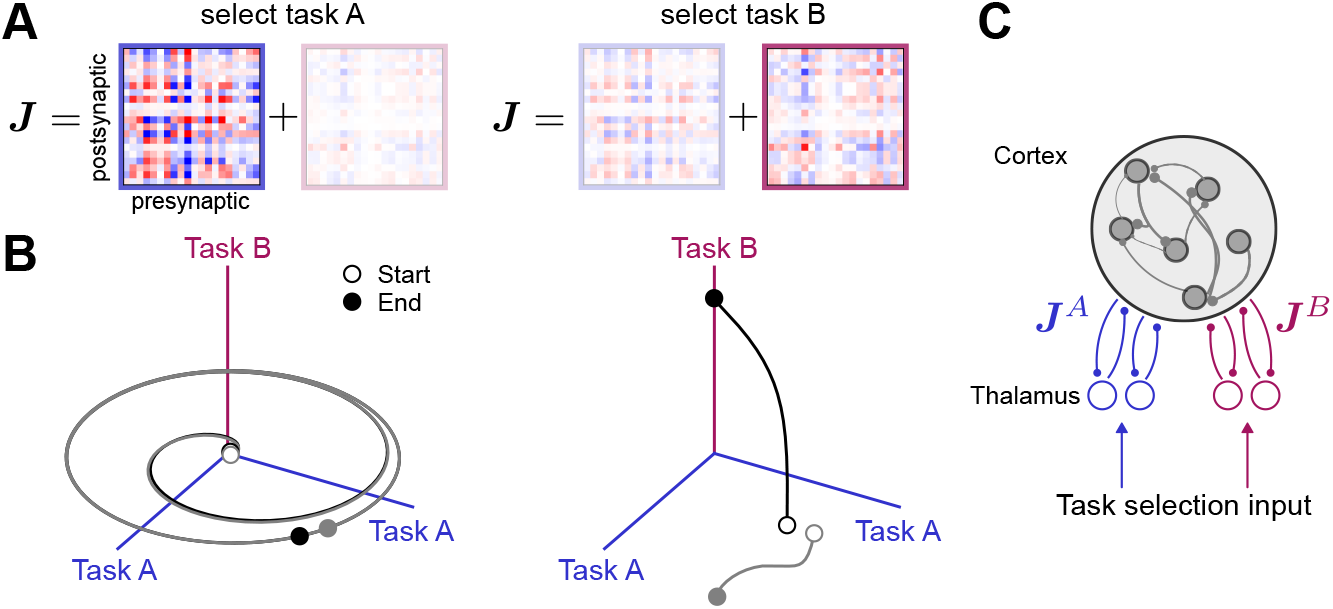
Sequential multi-tasking through changes in effective connectivity. **A** A model of the limit cycle and memory tasks with the same connectivity as in Fig. 3 and an additional factor that increases the magnitude of the task A or B connectivity component when the respective task is selected. **B** Two example neural trajectories (gray and black) over sequential selection of task A (left) and B (right). In contrast to Fig. 3, initial conditions are not precisely aligned to each subspace. Initial conditions for the right plot are endpoints of trajectories for the left plot. When task B is selected, an input, representing a sensory cue and either positively (black) or negatively (gray) aligned with the task B subspace, is briefly provided to perturb the neural trajectories toward different fixed points. **C** Anatomical basis for modulation of effective connectivity proposed by Logiaco et al. ^15^. Connectivity components are associated with distinct corticothalamic loops.

An extreme version of this mechanism that sets *a*_*µ*_ = 1 for a selected task and *a*_*ν*≠*µ*_ = 0 for all other tasks enables sequential multi-tasking by completely overwriting the synaptic weight matrix each time a new task is selected. Several models make this assumption^15,18^. However, recent work has shown that such dramatic alterations in weights are not required in nonlinear networks ^30^. Instead, to select task *μ*, modestly increasing only *a*_*µ*_ suffices. The amount of increase required depends on the distribution of *a*_*ν*≠*µ*_ as well as the number of tasks *P* relative to the network size *N*. Thus, nonlinear cross-task interference aids rather than hinders multi-tasking by allowing the selected task’s dynamics to suppress those of unselected tasks.

When a task is selected through this mechanism, the associated manifold is globally attracting. This permits multi-tasking through sequential modulation of the *a*_*µ*_ associated with different tasks. Fig. 5B illustrates selection of task A (left) followed by task B (right) by increases in *a*_*A*_ and *a*_*B*_. However, such a mechanism alone does not enable simultaneous multi-tasking.

What biophysical processes may support task-related changes in effective connectivity? Previous studies have suggested that neuromodulation^43^ or short-term synaptic plasticity may modify the effective weight between pairs of neurons, although ensuring that these modifications are specific to individual connectivity components may be challenging^30^. An attractive interpretation associates each connectivity component with distinct corticothalamic loops whose gains are selectively modulated, for instance by the activity of basal ganglia output nuclei, to select a task^15^ (Fig. 5C).

#### Selection by a rule input

Transient activity or modulation of effective connectivity underlie many mathematical models of multi-task activity ^12,15,30,35^. However, recurrent neural networks trained to perform flexible computations through gradient descent typically rely on a different mechanism. A tonic external input, sometimes called a “rule” input, excites or inhibits a task-specific pattern of neurons and switches each time a new task is selected^9,10^ (Fig. 6A). Like in the previous section, this implies that the flow field changes upon task selection, but this change is not due to direct modulation of the weight matrix but rather the nonlinear effect of the rule input.

**Figure 6.**
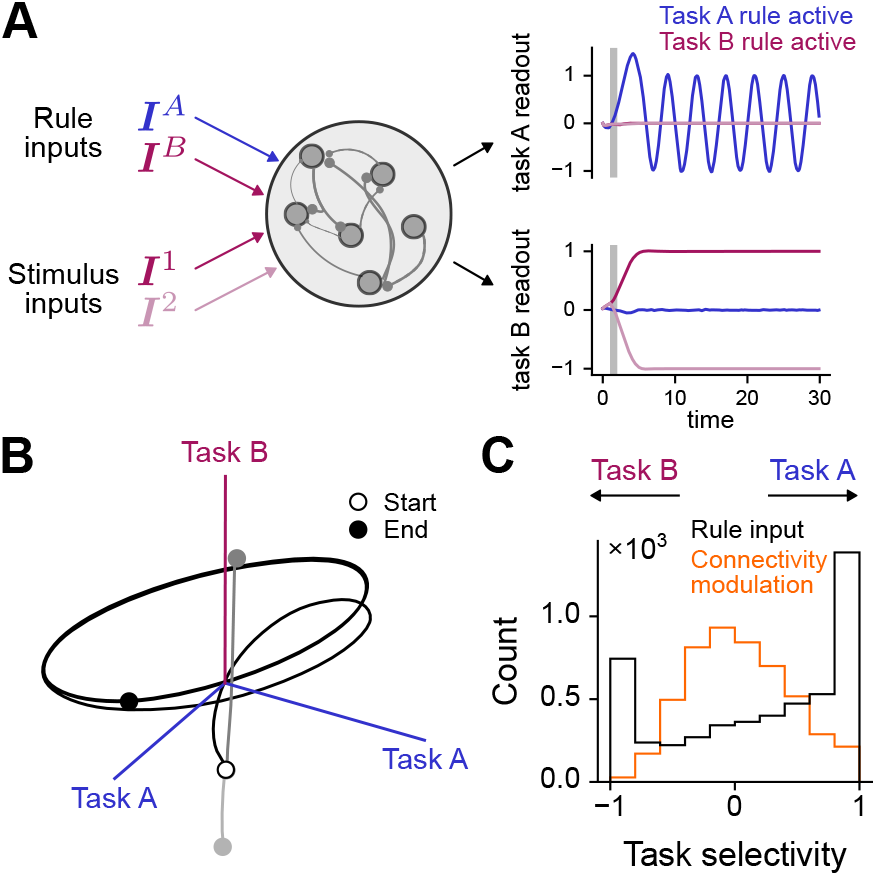
Sequential task selection by a rule input, in a full-rank network trained on the limit cycle and memory tasks introduced in Fig. 3. **A** Schematic illustrating a network receiving rule inputs to select task A or B and stimulus inputs to select the task B fixed point. Right: Linear readouts of a network trained to produce an oscillation when the task A rule input is active and bistable fixed points when the task B rule input is active. Loss is determined by readout activity for *t >* 5. When task B is selected, a stimulus input indicating the correct task B fixed point is provided during the time interval indicated by the gray box. **B** Example neural trajectories with the same initial condition during selection by task A (black) or B (gray) rule inputs. Axes are defined by projections onto the top two principal components of task A trials and the top principal component of task B trials. Light and dark gray trajectories represent trials with distinct stimulus inputs. **C** Distribution of neuronal task selectivity index (*σ*_*A*_ − *σ*_*B*_)/(*σ*_*A*_ +*σ*_*B*_), where *σ*_*µ*_ is the standard deviation of a neuron’s activity during the performance of task *μ*. Distribution is computed across neurons of 10 trained networks, each with *N* = 500 (black), and compared to the distribution for networks that multi-task through modulation of effective connectivity as described in Fig. 5 (orange).

A recurrent neural network trained in this manner, without a constraint on the rank of its weight matrix, successfully performs the limit cycle and memory tasks examined in Figs. 3–5. Neural activity projected onto subspaces defined by the top principal components of activity during performance of each task follows trajectories similar to those of previous mechanisms (Fig. 6B). These task subspaces are nearly orthogonal (mean subspace angle of 80 *±* 6 degrees across 10 trained networks) and account for a large fraction of the variance of neural activity during performance of each task (0.85 ± 0.1 for task A and 0.994 ± 0.001 for task B). These properties of orthogonality and low dimensionality of task subspaces in trained networks are similar to those of the low-rank networks we have examined so far, although the training procedure did not explicitly require low-rank weight updates or low-dimensional activity.

However, the distribution of individual neuron selectivities to the two tasks differed from previous mechanisms. We computed a task selectivity index ranging from −1 (purely modulated by task B) to 1 (purely modulated by task A). Neurons exhibited a range of selectivities, but a much larger fraction had values near the extremes as compared to, for example, networks that multi-task through modulation of effective connectivity (Fig. 6C). This observation is consistent with prior studies that report the emergence of task-specific modules of neurons over gradient descent training^9,10^.

#### Barcoded connectivity as a solvable model of multi-tasking

Although rule inputs thus support multi-tasking in networks trained through gradient descent, an analytically tractable model of this task selection mechanism is not yet available (Supp. Fig. S1; see Appendix). We therefore developed a new multi-task network model, motivated by the high selectivity of neurons in trained networks (Fig. 6C) and the fact that recurrent neural networks with groups of neurons whose connectivity statistics differ are more flexible in producing time-varying activity ^16,21^. In the model, a quantity 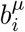 describes the degree to which neuron *i*’s connections are tuned to task *μ* (Fig. 7A, top). We call the vector ***b***^*µ*^ the task *μ* “barcode.” The connectivity vectors ***m***^*µ,r*^ and ***n***^*µ,r*^ in Eq. 3 are then defined as

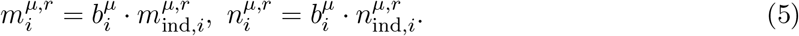

**Figure 7:**
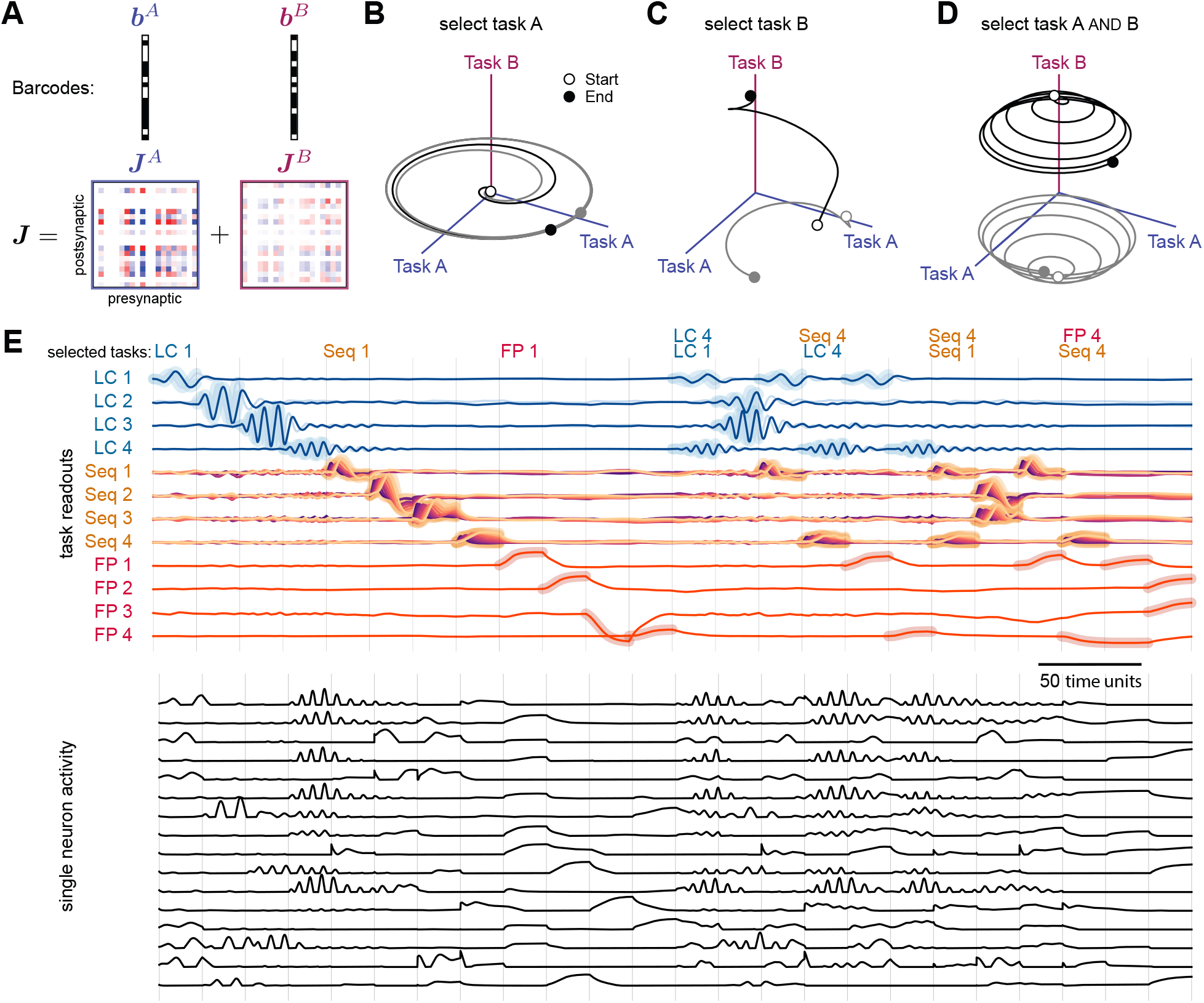
Barcoded connectivity. **A** Schematic of barcodes ***b***^*µ*^ (top; black squares indicate positive values) and corresponding connectivity components ***J*** ^*µ*^ (bottom). **B** Two example neural trajectories (gray and black) during selection of task A by a rule input. **C** Similar to **B** but during selection of task B. As in Fig. 5B, an input positively (black) or negatively (gray) aligned with the task B subspace is provided when task B is selected. **D** Similar to **C** but during simultaneous selection of both tasks by a rule input aligned with both barcodes. **E** Example network that performs 12 tasks: four limit cycles (LC), four sequences (Seq) and four fixed point attractors (FP). Different combinations of tasks are selected in each time epoch. Gray lines denote epoch boundaries. Top: Linear readouts of the different task manifolds. The highlighted background indicates when the corresponding task is selected by the rule input (labels at top of panel also indicate the selected task; not all epochs are labeled). For the first 12 epochs, only single tasks are selected, while for the last 12 epochs, two tasks are simultaneously selected. Bottom: Firing rates of 16 example neurons. Barcode sparsity parameter *f* = 0.3.

The elements of 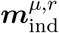 and 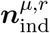 are random variables drawn independently for each neuron, as in previous sections. Unlike before, these elements are then multiplied by the barcode to generate the connectivity component for task *μ* (Eq. 3). To begin, we will assume that the barcode ***b***^*µ*^ is a binary vector in which a random fraction *f* of elements, chosen independently for each task, are nonzero (we generalize this in the Appendix). Each connectivity component ***J*** ^*µ*^ is additionally scaled by 1/*f*, which ensures that single-task dynamics are preserved across *f*. In this model, the barcode can be interpreted as defining those neurons whose synapses are modified so that the network can perform task *μ* (Fig. 7A, bottom). Because a neuron’s selectivity to each task is drawn independently, this model does not imply the existence of disjoint clusters of similarly selective neurons.

The introduction of barcodes enables rule inputs that are aligned to each task’s barcode to switch a network’s activity between task manifolds (Fig. 7B,C). A mathematical analysis demonstrates that this ability is due to the rule input selectively modulating the gain of the neuronal subpopulation defined by the selected task’s barcode (see Appendix). It is also possible to provide a rule input that selects multiple tasks at once if it is aligned with multiple barcodes, enabling simultaneous multi-tasking (Fig. 7D; see Appendix). In this case, neural trajectories both oscillate in the task A subspace and maintain a separation along the task B subspace, although the magnitude of each projection is moderately reduced compared to the single-task case. Thus, a single network may either sequentially or simultaneously multi-task, a flexibility that, to our knowledge, has not been explored in previous models.

One concern is that, if *f* is small (relative to 1*/P*), the model approaches the unrealistic extreme of disjoint task-specific populations that we rejected earlier. However, our theoretical analysis shows that very low values are not required. In fact, a fixed value of *f*, that does not decrease as the number of tasks *P* grows, is sufficient (see Appendix). This means that, in networks that support many tasks, individual neurons each participate in many tasks. As an example, we constructed a network with *f* = 0.3 that multi-tasks between 12 tasks that each require oscillatory, sequential, or fixed-point dynamics (Fig. 7E, top). Application of rule inputs, aligned either with a single barcode or multiple, enables sequential multi-tasking of individual tasks or simultaneous multi-tasking, respectively. Neuronal firing rates are heterogeneous and cannot be clustered into discrete response profile categories (Fig. 7E, bottom). The model therefore demonstrates that increased task selectivity, and the additional flexibility it provides for generating dynamics, does not require disjoint task-specific subpopulations of neurons.

## Discussion

### Comparison of multi-tasking mechanisms

Table 1 summarizes the four multi-tasking mechanisms we have described. Although new mechanisms may be introduced, categorical distinctions between those we have described are revealed. The first concerns the capacity for simultaneous multi-tasking, which is only enabled by rule inputs in the models we have considered (Table 1, second column). Another concerns the nature of the flow field that determines the neural dynamics (Table 1, third column). If tasks are associated with either transient or multistable activity, this flow field is unchanged across tasks, meaning that task selection must be achieved through a preparatory process that determines the initial state. Moreover, only in the case of multistability are multiple stable attractors corresponding to different tasks present at once. On the other hand, task selection through a change in effective connectivity or a rule input implies that the flow field is modified upon task selection. We note that these mechanisms are not mutually exclusive. For example, barcoded connectivity could enable modulation of effective connectivity to support simultaneous multi-tasking, or reduce the precision of initial condition required for transient task selection.

**Table 1.** Comparison of multi-tasking mechanisms.

| Task selection mechanism | Simultaneous or sequential | Dynamical picture | Control parameter | References |
| --- | --- | --- | --- | --- |
| Preparatory activity/transients | Sequential | One flow field, trajectory determined by initial state | Precision of initial state | 13, 35 |
| Multistability | Sequential | One flow field with multiple attractors, trajectory determined by initial state | Distribution of task selectivities | 29, 36–38 |
| Change in effective connectivity | Sequential | Flow field changes as connectivity changes | Magnitude of connectivity modulation ($a_\mu$) | 15, 18, 30, 43 |
| Rule input | Both | Flow field changes with rule input | Degree of modularity (barcode sparsity $f$) | 9, 10 |

In many of the models we examined, the mechanism that supports multi-tasking is associated with certain parameters that must be adjusted to combat the problem of interference (Table 1, fourth column). For task selection associated with transient activity, this parameter is the precision of the initial state. For modifications of effective connectivity, it is the magnitude of change in the connectivity component associated with the selected task. For models with barcoded connectivity, it is the degree of task selectivity, as measured in our model by the barcode sparsity *f*. Our mathematical analysis shows that, as task diversity (as quantified by the degrees of amplification associated with each task) and thus the potential for interference increases, each of these parameters must be adjusted more to compensate (see Appendix).

### Model simplifications and extensions

We have made several simplifying assumptions to derive our theory. One is to assume that each task corresponds to an independent computation and is associated with an orthogonal subspace in neural firing rate space. This assumption is made by many theoretical models ^13,15,18,30^. Experiments with multiple task epochs, such as fixation, delay, and response periods, may be best thought of as a series of tasks, each corresponding to a single such epoch, in our model ^18^. However, certain tasks may instead be described as parametrically related to one or multiple others, for instance adjusting the speed or timing of a motor program^44^. In this case, one may expect the overlap between task subspaces to smoothly depend on this parameter^45^. Alternatively, such situations may be conceived as a single, higher-dimensional task in which the parameter is identified with a new task variable.

We also focused on neural networks that generate time-varying, nonlinear dynamics, which lead to the phenomenon of interference that our theory examines (Fig. 2D). Neurons themselves are nonlinear, and restricting to linear dynamics greatly limits the range of computations that can be performed, arguing against models of neural dynamics that are exclusively linear. However, it should be noted that for brain regions that primarily respond to external inputs from other regions, rather than generating sustained patterns of activity, amplification that leads to task interference may be weak. We therefore predict that the mechanisms we describe are particularly important for brain regions that support working memory and motor pattern generation.

For simplicity, most of our examples have illustrated multi-tasking with two tasks. Our theory generalizes to any finite number of tasks *P*, in the limit *N* ≫ *P* (see Appendix). When *P* is comparable to *N*, more analysis is required, and theoretical results exist only for certain cases. These include analyses of capacity for storage of fixed-point or continuous attractors^31,37,46^. For storage of attractors or sequences with asymmetric connectivity, the limit of large *P* may be associated with the emergence of chaotic activity due to interference between tasks^13,47^. A similar phenomenon was analyzed in a recent study that focused on multi-tasking through changes in effective connectivity^30^. Defining and calculating a general notion of capacity for multi-tasking recurrent neural networks is an open problem.

Our analysis focuses on dynamics on task manifolds while remaining agnostic as to the readout of these dynamics. Distinct tasks may be associated with distinct linear readouts, or the weights of a common readout may overlap with multiple task manifolds. Alternatively, readout-related activity may correspond to a dimension in neural activity space that is shared across tasks but does not interfere with each task’s recurrent dynamics (see Appendix).

As shown in the Appendix, only one task is transversely stable and all the rest are unstable if the elements of the connectivity vectors, 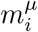 and 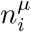, are drawn independently from a jointly Gaussian distribution, as in many low-rank network models^20,30,48^. Task barcodes (Fig. 7) are a parsimonious relaxation of this Gaussian assumption and consistent with observations of modular neural subpopulations in experimental recordings^49,50^ and task-trained networks^9,10^. In models of continual learning, barcodes may be produced by a task-specific signal that selects a random subset of neurons for synaptic weight changes and does not require pre-allocation of neuronal subpopulations for distinct tasks. However, other non-Gaussian distributions of connectivity vector elements, for example cosine-tuned connectivity vectors^29,38^ (Fig. 4), can also support multi-tasking. We provide a general mathematical analysis of the effects of these distributions on multi-tasking in the Appendix.

### Sequential and simultaneous multi-tasking in experiments

Forms of sequential multi-tasking have been examined in various experimental paradigms. These include the production of different movement patterns in trained visuomotor tasks in primates^51–53^ and rodents^54^. Natural behavior is often viewed as a sequence of distinct motor patterns^4,55–58^. Many of these studies report the existence of distinct task subspaces^4,51,53,58^ and high neuronal task selectivity^54,57^, consistent with our models.

Although naturalistic behavior often involves the generation of distinct movement patterns across different effectors, simultaneous multi-tasking has been studied relatively less frequently in experiments. The most common examples are forms of memory-guided tasks in which multiple items must be stored in memory at once. Recent studies have suggested that, in such situations, each item may be stored in a largely orthogonal subspace, or “slot” ^59–62^. Executing multiple tasks at once often degrades performance, a phenomenon known as dual-task interference^63^. Recordings in primate lateral prefrontal cortex demonstrate that this interference is associated with neurons that participate in both tasks^24^, consistent with our analysis which shows that high single-neuron task selectivity supports simultaneous multi-tasking (Fig. 7).

A system capable of sequential or simultaneous multi-tasking can be viewed as a compositional system, in that it can perform computations in different orders or combinations^64^. Indeed, trained animal behaviors that involve sequential^51–53^ or simultaneous^65^ multi-tasking are sometimes described as compositional. We have focused on how a single neural network performs multiple distinct computations through a selection mechanism, which is a prerequisite for compositional behaviors unless each computation is associated with a dedicated subset of neurons. A complete understanding of such behaviors will also require an understanding of the systems that represent rules dictating which computations should be selected and when. Some forms of composition may also involve rules for combining computations that extend beyond sequential or simultaneous multi-tasking^64,66^.

### Signatures of multi-tasking mechanisms in neural data

In experiments with explicit task structure, task subspaces may be identified by performing dimensionality reduction on independent subsets of data. An unsupervised method for identifying putative task-dependent factors was recently proposed that may be applied to experiments without such explicit structure^67^. Several studies have quantified the overlap of identified task subspaces, reporting increased alignment for more similar tasks. This may reflect similarity of the stimulus features^53^, sensorimotor transformations^45,51^, or muscle groups^68^ required for task performance. The degree of subspace alignment varies across brain regions: sensory areas tend to reuse subspaces across tasks, while cognitive and motor regions tend to orthogonalize ^4^. In higher regions, certain task dimensions have been identified as responsible for executing components of a given task that may be shared with other tasks or task conditions^53^. A recent study suggests that motor cortex may combine “subskills” whose associated motor readouts additively produce a desired trajectory, a form of simultaneous multi-tasking if each subskill is identified as a single task^51^. Activity across epochs each requiring multiple such subskills may therefore span non-orthogonal subspaces^45,51,68^.

Methods for inferring the flow field associated with neural dynamics have been proposed^69^, some of which involve fitting low-rank recurrent neural networks to data^70^. Evaluating whether the flow field is task-dependent, in regions of neural state space that are explored in multiple tasks, will distinguish mechanisms that rely on initial conditions or changes in the flow field for task selection. Optogenetic perturbations that disrupt preparatory activity and bias activity away from one task manifold and toward another may also be used to probe this distinction. O’Shea et al. ^71^ studied the effects of optogenetic and electrical microstimulation perturbations to activity in primate motor cortex during a reaching task. This revealed activity consistent with a low-dimensional, transversely stable subspace. This provides evidence against the transient mechanism we described, though it does not immediately distinguish between other mechanisms.

The degree to which individual neurons are tuned to specific task variables or exhibit mixed selectivity to all variables has been debated^34,50,72,73^. Analyses of feedforward networks have demonstrated that mixed selectivity to multiple task variables increases the dimension of a neural representation and the linear discriminability of distinct stimulus classes ^34^. Analysis of neuronal selectivity across many mouse brain areas found that such mixing is common and did not identify evidence for clustering into discrete functional neuron types^74^. In contrast, analyses of recurrent models show that subpopulations of neurons with structured selectivity to distinct variables increase the flexibility with which a network can generate dynamics ^16,21^.

Our model of task barcodes suggests that when an individual network is responsible for generating very different dynamics, the required degree of task selectivity increases. It also suggests that neuronal populations selective to tasks that are more prone to interference may have reduced overlap. In this model, task variable encoding is distributed across neurons but the degree of selectivity is continuously parameterized by the barcode sparsity. Thus, the model permits interpolation between the extremes of disjoint task-specific subpopulations and fully distributed representations. Moreover, because a neuron’s selectivity to each task is drawn independently, the model does not predict clusters of neurons with identical selectivities. More broadly, our analysis identifies the distribution of task selectivities across a neural population as an important determinant of its multi-tasking capability.

## Acknowledgments

M.B., O.M., K.Z., and A.L.-K. were supported by the Gatsby Charitable Foundation (GAT3708). M.B. and A.L.-K. were also supported by NIH award RF1DA060772. M.B. was also supported by NSF DBI-2229929. K.Z. was also supported by NIH award 1F32NS146149. A.D. and S.O. were supported by the CRCNS project PIND (ANR-19-NEUC-0001-01), the Simons Foundation (AN-NC-GB-Culmination-00003154-05, SO), the European Union (ERC Synergy project CHRONOLOGY, grant agreement No 101167367) and the program “Ecoles Universitaires de Recherche” (ANR-17-EURE-0017).

## Methods

### Network dynamics

We consider a recurrent neural network of *N* firing rate neurons, whose dynamics and connectivity are defined by Eqs. (1) and (3). A low-dimensional description of the network dynamics can be obtained by projecting neural activity *x*_*i*_ onto the connectivity vectors associated with each task and the mean activity direction:

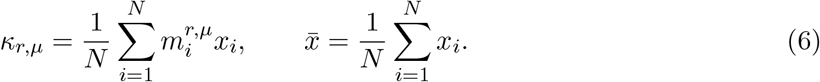

In the Appendix, we analyze the dynamics of these projections in the limit of large *N* using the theory of low-rank recurrent neural networks.

### Task-related connectivity

We generate task-related connectivity components by specifying the distributions of the connectivity vector elements 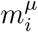 and 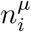 in Eq. (2). For the bistable working-memory task, we use a rank-one component:

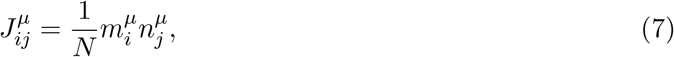

where 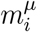 and 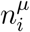 are sampled from a Gaussian distribution with zero mean and covariance *σ*_*µ*_ unless otherwise specified.

For the generation of limit cycle dynamics, we use a rank-two component

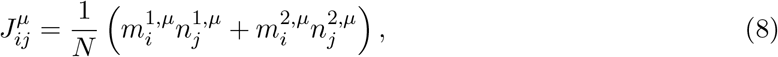

where the elements of the four connectivity vectors 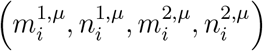 are drawn from a four-dimensional Gaussian distribution, unless otherwise specified, with zero mean and covariance 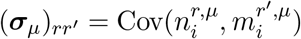

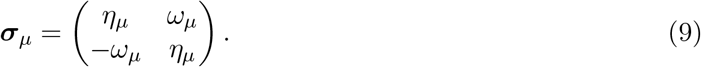

All other pairs of connectivity vector elements, including those of different tasks, are uncorrelated. For each task, the variance of 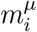 is set to one and the variance of 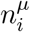 is set to a constant such that the covariance matrix defining the distribution of all connectivity vector elements is positive definite. Note that the value of this constant does not affect the dynamics in the large *N* limit.

In Fig. 2, we study the dynamics of a network with two connectivity components each associated with limit cycle dynamics. The task parameters are *η*_*A*_ = 1.3, *ω*_*A*_ = 0.45 for task A and *η*_*B*_ = 1.7, *ω*_*B*_ = 1.7 for task B. The network has *N* = 1000 neurons with a rectified-linear activation function *ϕ*(*x*) = max(*x* −*θ*, 0) with threshold *θ* = −1. The task outputs are projections of the firing rates *ϕ*(***x***) onto the readout vectors ***m***^1,*A*^/(0.8*N*) and ***m***^1,*B*^/(3*N*), respectively.

### Models of task selection

#### Transient activity

The network of Fig. 3B,C contains *N* = 10,000 neurons with rectified-linear activation functions. The threshold is *θ* = −1, and *g*_*I*_ = 2. Task A is associated with limit cycle dynamics with *η*_*A*_ = 1.6, *ω*_*A*_ = 0.8, and task B is associated with bistable dynamics with *σ*_*B*_ = 1.8. Task selection is determined by the initial condition. For task B, the initial condition is 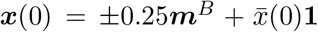, where 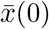 is determined as the self-consistent solution to 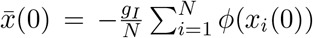. For task A, it is 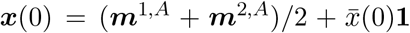. Trajectories correspond to 60 time units.

The network of Fig. 3D is similar to the above except *σ*_*B*_ = 1.7. The initial conditions are 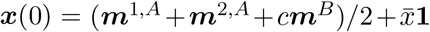, where *c* determines the task B projection. The simulation is repeated for *c* = ±|*c*|, and the reported duration is the minimum across these two repetitions.

#### Multistability

In Fig. 4A,B, we study a network with two connectivity components supporting ring-attractor dynamics, with *N* = 10,000 neurons with rectified-linear activation functions and *θ* = −1. For each task, preferred angles 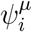 are assigned to each neuron by randomly permuting equally spaced angles on [0, 2*π*). The synaptic weight matrix is

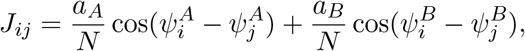

with *a*_*A*_ = 2.5 and *a*_*B*_ = 2.3. Thus, the connectivity patterns are cosine- and sine-distributed.

In Fig. 4C,D, a rank-two limit cycle component is combined with a rank-one component supporting fixed points in a network of *N* = 1000 neurons with the same activation function. The synaptic weight matrix is

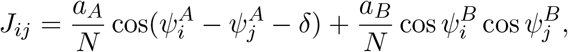

with *δ* = 0.35*π, a*_*A*_ = 2.2/(cos *δ*), and *a*_*B*_ = 2.6. The two trajectories share an initial condition and receive opposite instantaneous perturbations Δ***x*** = ±1.2***m***_*B*_ at *t* ≃ 19.8, where 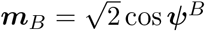. Readouts are projections of *ϕ*(***x***) onto the corresponding task patterns, normalized by *N*. The task Apattern is defined as 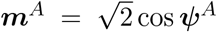, while the task B pattern was first defined as 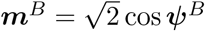 and was subsequently orthogonalized with respect to patterns cos ***ψ***^*A*^ and sin ***ψ***^*A*^. The perturbation was also provided along this orthogonalized B pattern.

#### Modulation of effective connectivity

The parameters of the network in Fig. 5 are the same as those of Fig. 3. Task selection is achieved by setting *a*_*µ*_ = 1 for the selected task and *a*_*µ*_ = 0.5 for the unselected task. The initial condition *x*_*i*_(0) when task A is selected is drawn independently from a standard Gaussian distribution. When task B is selected, for the first 10 time units, an external input 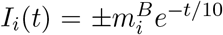 perturbs the trajectory toward one of the two fixed points. Trajectories correspond to 60 time units.

#### Task-trained network with rule inputs

The network of Fig. 6 contains *N* = 500 neurons with rectified-linear activation functions and thresholds *θ*_*i*_. Activity evolves according to:

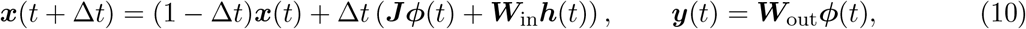

where the length-4 vector ***h***(*t*) denotes external input channels, ***W***_in_ is the *N* × 4 input weight matrix, ***W***_out_ is the 2 × *N* output weight matrix, ***y*** = (*y*_*A*_, *y*_*B*_)^⊤^ denotes linear readouts for tasks A and B, and Δ*t* = 0.1. The elements of each weight matrix are initialized as independent Gaussian variables with standard deviations 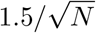 for ***J***, 0.5 for ***W***_in_, and 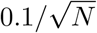 for ***W***_out_. Thresholds are initialized as *θ*_*i*_ = −0.2 for each neuron.

The four input channels ***h*** = (*h*_*A*_, *h*_*B*_, *h*_1_, *h*_2_) consist of two binary rule channels *h*_*A*_ and *h*_*B*_ that are nonzero when task A or B is selected and two binary stimulus channels *h*_1_ and *h*_2_, one of which is nonzero during the interval *t* ∈ [1, 2) for task B trials. Inputs depicted in Fig. 6A are determined by the columns of ***W***_in_***h***. Each trial is of length 60 time units, and ***x***(0) = **0**. For task A trials, the target output is 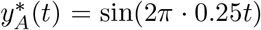. For task B trials, the target output is 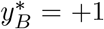 if the network receives input *h*_1_ or 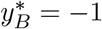 if the network receives input *h*_2_.

Networks are trained using backpropagation through time and the Adam optimizer to minimize the mean squared difference between *y*_*µ*_(*t*) and 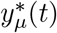 computed over the interval *t* > 5, where *μ* denotes the current trial’s task. All parameters ***J***, ***W***_in_, ***θ***, and ***W***_out_ are optimized. Each training batch contains 3 trials, one each of task A, task B with input 1, and task B with input 2. During training, the learning rate exponentially decays from 10^−3^ to 10^−4^ over 1000 epochs, and gradients are clipped to a maximum norm of 1.

#### Barcodes

The network of Fig. 7B–D is similar to those of Figs. 3 and 5, except that the connectivity is constructed using the barcode model (Eq. 5) with *f* = 0.7, *η*_*A*_ = 1.75, *ω*_*A*_ = 1, and *σ*_*B*_ = 1.7. The input to the network is 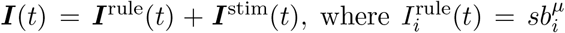 to select task *μ* and 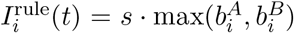 to select both tasks simultaneously, and *s* = 2. As in Fig. 5, the initial condition *x*_*i*_(0) when task A is selected is drawn independently across neurons from a standard Gaussian distribution, and when task B is selected, for the first 10 time units, a stimulus input 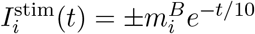 perturbs the trajectory toward one of the two fixed points. Trajectories correspond to 60 time units.

The network of Fig. 7E multi-tasks with four limit cycle tasks, four feedforward sequences, and four fixed-point tasks in a network of 6000 rectified-linear neurons. The threshold *θ* = −0.3, and *g*_*I*_ = 1.6. Each task is associated with a non-binary barcode (see Appendix A.8.3). A fraction of exactly *f* = 0.3 of barcode entries are set to one. The other entries are zero for the ***n***^*µ*^ patterns and *δ* = 0.1 for the ***m***^*µ*^ patterns. The limit cycle parameters (*η*_*µ*_, *ω*_*µ*_) are (1.5, 0.6), (1.55, 1.0), (1.6, 1.5), and (1.65, 1.8). For the sequential tasks (Eq. 4), the number of dimensions *R*_*µ*_ per task and the feedforward gain (*R*_*µ*_, *g*_*µ*_) are (8, 1.4), (10, 1.45), (12, 1.5), and (14, 1.55). For the fixed-point tasks, strengths *σ*_*µ*_ are 1.5, 1.6, 1.65, and 1.7. Tasks are selected individually or in pairs in epochs of length 21 time units by applying an input of amplitude *s* = 0.6 to neurons in the union of the selected barcodes. When a sequence is selected, the network receives an input proportional to the first pattern, divided by 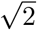 when two sequences are selected simultaneously.

**Figure S1:**
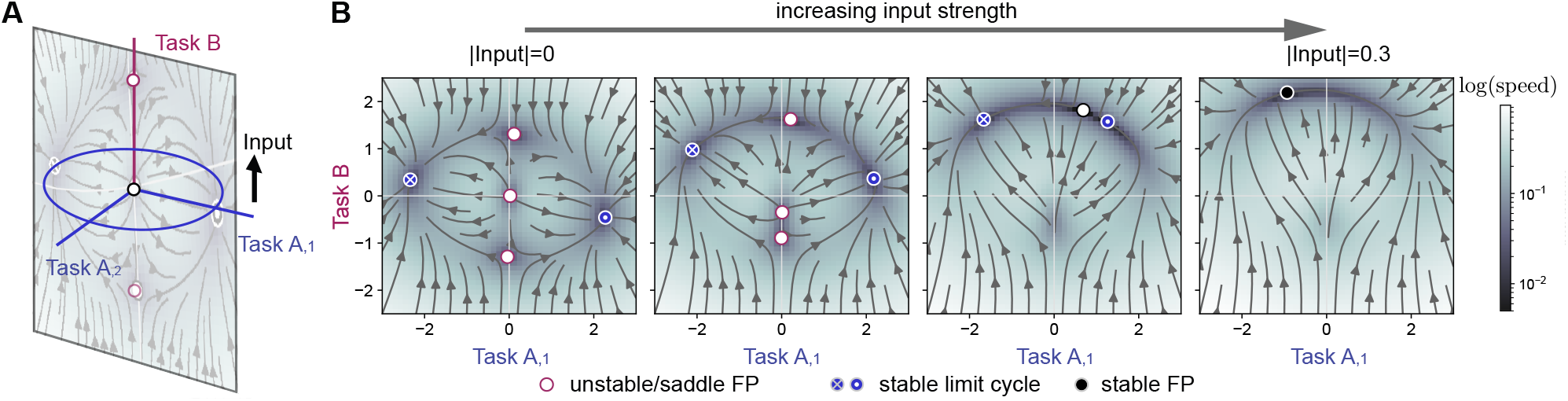
External inputs with Gaussian statistics do not affect relative transverse instability. **A** Illustration of a rank-three network with Gaussian connectivity statistics, performing task A (a limit cycle, blue line) and task B (bistable fixed point). Task B is unstable when perturbed along the Task A dimension. We add a tonic Gaussian input along the Task B axis (black line, input parallel to ***m***^*B*^). We focus on the dynamics on the illustrated two-dimensional slice. **B** Flow field along the two-dimensional plane defined by the task-B variable and one of the two task-A variables. The different panels show increasing input strength (0, 0.1, 0.2 and 0.3). The input along Task B cannot stabilize Task B. As the input increases, the network undergoes a first bifurcation, where the bistable fixed points along the task-B manifold turn into a single input-driven fixed point, which remains transversely unstable. A second bifurcation happens when the stable limit cycle collapses into an input-driven stable fixed point. The recurrent dynamics associated with task B are never stable, independently of the external input. Network size *N* = 10,000.

**Figure S2:**
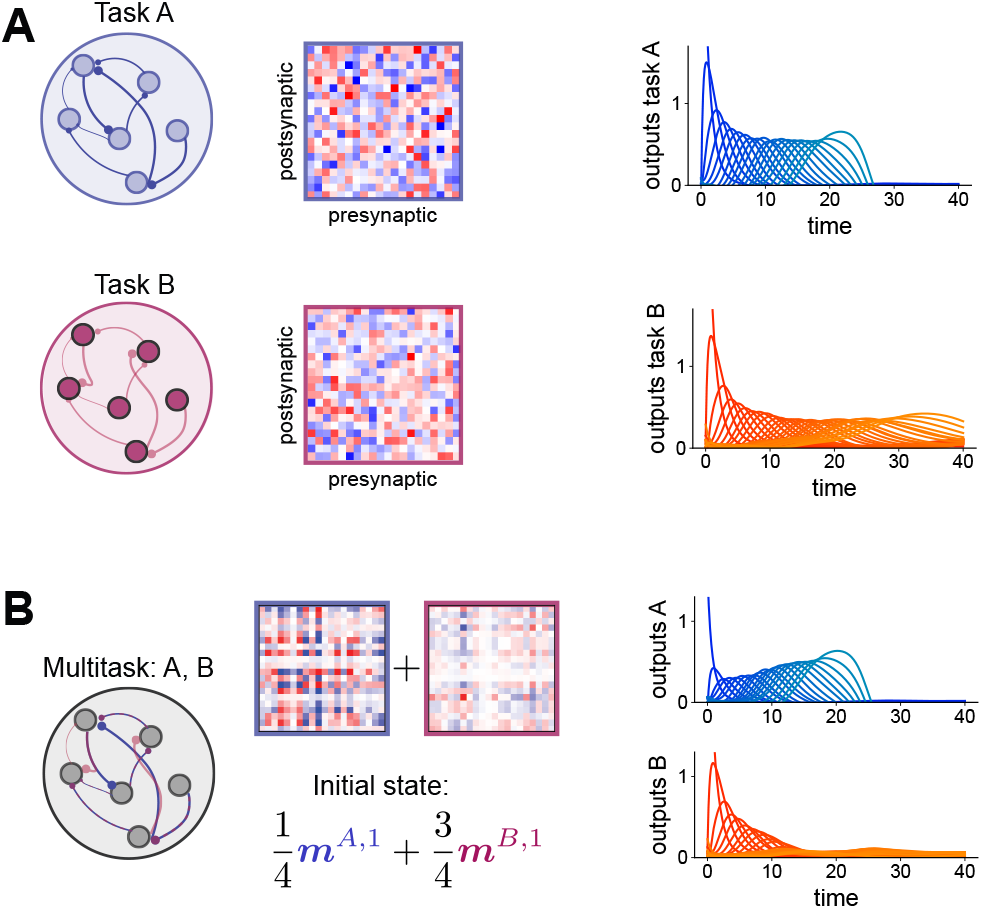
Interference between sequences. **A** Task A consists of a sequence of 20 different patterns, Task B consists of a sequence of 30 different patterns. Individual networks can reproduce such sequence of activity, with a sequence duration approximately equal to the number of patterns. The connectivity follows Eq. (4) ^13^. **B** When the connectivity matrices corresponding to both tasks are combined into a single network, the task with the strongest weights (task A in this example) is stable, while task B becomes transversely unstable. When initialized with an initial state overlapping with both sequences, only task A can be executed until the end, even if task B has a stronger initial overlap.

## Appendix Supplemental theoretical results

### A.1 Definition of multi-tasking manifolds

Consider a set of tasks indexed by *μ* = 1 … *P*, each defined by dynamics *F*_*µ*_ on its task variables ***κ***_*µ*_:

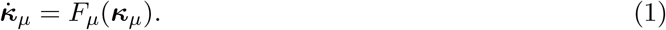

The vector ***κ***_*µ*_ has *R*_*µ*_ elements and represents quantities relevant for performance of task *μ*.

We now consider a recurrent neural network and define what it means for the dynamical system governing its activity to have manifolds that permit sequential or simultaneous multi-tasking of tasks defined by Eq. (1). Suppose there is an invariant manifold in the network’s activity space, and locations on the manifold are smoothly parameterized by ***z***. Denote the dynamics of ***z*** on this manifold by ***ż***= *G*(***z***). The function *G* is determined by the dynamical system that governs neural activity, evaluated at locations on the manifold.

We next need a notion of what it means for the dynamics of ***z*** to be equivalent to those of the task variables (Eq. 1). We use a property called *topological conjugacy*, which, informally, requires existence of a continuous, invertible transformation that maps trajectories of one dynamical system to another while preserving time. This notion has been previously used to assess similarity of dynamics in neural activity space ^69^. Conjugate dynamical systems have the same global structure, such as their arrangements of fixed points and limit cycles. Formally, if two dynamical systems are defined by^†^ ***z***(*t*+*T*) = *ϕ*_*T*_ (***z***(*t*)) and ***κ***(*t*+*T*) = *ψ*_*T*_ (***κ***(*t*)), they are topologically conjugate if there is a homeomorphism ***κ*** = *h*(***z***) such that *h* ∘ *ϕ* = *ψ* ∘ *h*. This notion of equivalence is intentionally broad, and for several of the examples we consider *h* is the identity function.

With this definition, we say that a system *supports sequential multi-tasking of tasks μ and ν* if it has a distinct attracting manifold for each task and the dynamics on these manifolds are conjugate to those defined by *F*_*µ*_ and *F*_*ν*_, respectively. The dimensions of these manifolds are therefore *R*_*µ*_ and *R*_*ν*_.

To define dynamics associated with performing tasks *μ* and ν simultaneously, let ***κ*** = (***κ***_*µ*_, ***κ***_*ν*_) with dynamics given by 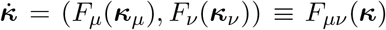. This dynamical system corresponds to two uncoupled sets of task variables, each following Eq. (1). A system *supports simultaneous multi-tasking of tasks μ and* ν if it has an attracting manifold and the dynamics on this manifold are conjugate to those defined by *F*_*µν*_. The dimension of this manifold is therefore *R*_*µ*_ + *R*_*ν*_. This definition can be generalized to more than two tasks by defining ***κ*** in terms of each task’s variables.

In some cases, there may be neural activity dynamics unrelated to the task variables, leading to higher-dimensional manifolds. In this case, the definitions can be extended by defining the dynamics in terms of ***κ*** = (***κ***_*µ*_, ***κ***_0_) or ***κ*** = (***κ***_*µ*_, ***κ***_*ν*_, ***κ***_0_) for sequential and simultaneous multi-tasking, respectively, where ***κ***_0_ describes these unrelated dynamics.

To show that a manifold satisfies one of these definitions, it is necessary to show both that it is attracting and that the dynamics on it are conjugate to those of the appropriate task variables. The networks we construct that perform transient multi-tasking (Fig. 3) satisfy the conjugacy requirement but not the attracting requirement. We note that, depending on which values of ***κ***_*µ*_ are relevant for task performance, one might choose to require these properties hold for only a subset of task variable values. For example, if ***κ***_*µ*_ represents a set of joint angles, the behavior of Eq. (1) for unachievable joint configurations is unimportant. We now investigate these properties for specific low-rank recurrent neural network models.

### A.2 Task variable dynamics in low-rank recurrent neural networks

We assume a network of the form

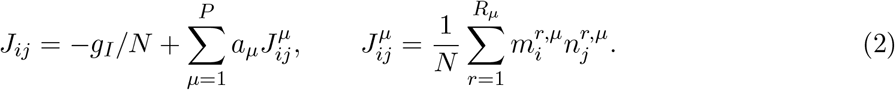

Except for the case of modulation of effective connectivity, (Section A.6.4), we assume *a*_*µ*_ = 1 for all *μ*. Activity in the network can be parameterized using a set of latent variables *κ*_*µ,r*_(*t*), along with a variable 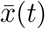 representing mean activity:

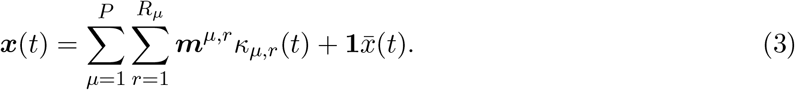

Activity decays along all directions except the span of {***m***^*µ,r*^} and **1**. For now, we further assume that there is no global inhibition, *g*_*I*_ = 0, such that 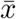 approaches 0. Later we will discuss how inhibition, and therefore time-varying values of 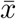, affects the dynamics (Sec. A.9). An advantage of considering low-rank recurrent neural networks is that, in terms of ***x***(*t*), task-related activity corresponds to a linear subspace rather than a nonlinear manifold (although the nonlinearity *ϕ*(***x***) produces a nonlinear manifold in firing rate space). We define the *task subspace* associated with task *μ* as the subspace of ***x*** in which ***κ***_*µ*_ takes any value, but ***κ***_*ν*_ = **0** for all *ν* ≠ *μ*.

The starting point for our analysis is the assumption that single connectivity components alone are sufficient to perform single tasks. That is, if 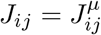, the variables ***κ***_*µ*_ that define the neural activity state (Eq. 3) follow the desired dynamics 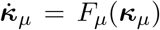. This assumption is based on extensive prior literature on low-rank recurrent neural networks as models of single-task computation^16,19–21^.

To simplify notation, we will describe most results in the setting of networks that perform two tasks *μ* ∈ *{A, B}*, but generalizations to more tasks will be noted. The dynamics are

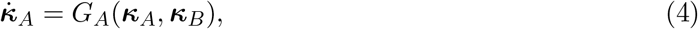

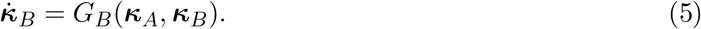

The key issue is that *G*_*A*_ and *G*_*B*_, in general, depend on both ***κ***_*A*_ and ***κ***_*B*_ and are thus different from *F*_*A*_ and *F*_*B*_. We refer to this coupling of task variables across different tasks as “interactions.” The form of these interactions determines whether a network with summed connectivity components satisfies the criteria for multi-tasking.

### A.3 Readouts of task-related activity

Our criteria for multi-tasking focus on dynamics on task manifolds without requiring the form of the readout of this activity to be specified. The simplest interpretation associates to each task a distinct linear readout *y*^*µ*^, defined by weights ***w***^*µ*^ that have nonzero alignment with the *μ*th task manifold, so that *y*^*µ*^(*t*) = ***w***^*µ*^ · *ϕ*(***x***(*t*)). Alternatively a shared readout *y*(*t*) may be generated through a single weight vector given by ∑_*µ*_ ***w***, so that *y*(*t*) = (∑_*µ*_ ***w***) · *ϕ*(***x***(*t*)).

It is also possible to modify the synaptic weight matrix, Eq. (2), so that readout-related activity is represented by the projection along a distinct subspace in neural activity space. Specifically, suppose

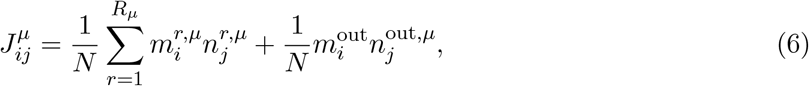

where the elements of ***m***^out^ are random and uncorrelated with all other connectivity vectors, and ***n***^out,*µ*^ is a vector in the *μ*th task subspace along which readout-related activity evolves. This creates an effectively feed-forward projection of neural activity along ***m***^out^, which forms a readout that is shared across tasks but only weakly feeds back into the task-related dynamics. These definitions can be straightforwardly generalized to multiple readout dimensions.

### A.4 Transverse stability

We require that a task manifold be attracting, meaning that nearby trajectories must return to it. A related requirement is what we will call *local transverse stability*, which is determined by the stability of the linearized dynamics orthogonal to the manifold and evaluated at a fixed location on the manifold. Local transverse stability of the *μ*th task subspace in networks of the form of Eq. (2) is determined by deviations in ***κ***_*ν*_, where *ν* ≠ *μ*, away from ***κ***_*ν*_ = **0**. These deviations can be characterized by the transverse Jacobian

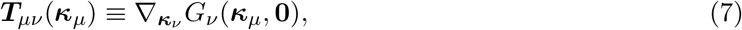

where 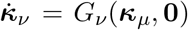 governs the dynamics of ***κ***_*ν*_ when ***κ***_*τ*_ = **0** for all *τ* ≠ *μ*. ***T***_*µν*_(***κ***_*µ*_) is thus interpreted as the linearization of the transverse dynamics of ***κ***_*ν*_, evaluated at a point ***κ***_*µ*_ in the task *μ* subspace. Because each subspace is orthogonal, local transverse stability can be evaluated by requiring, for each *ν* ≠ *μ* separately, that all eigenvalues of ***T***_*µν*_(***κ***_*µ*_) have negative real part.

Local transverse stability everywhere on a manifold does not always guarantee it is attracting^†^. However, it will be sufficient in our models because the transverse eigenvectors are the same at each point in a task subspace. If the associated eigenvalues always have negative real parts, projections onto these eigenvectors will decay.

### A.5 Simultaneous multi-tasking in linear networks with orthogonal subspaces

We first consider networks where the single-neuron transfer function is *ϕ*(*x*) = *x*. In this case, defining 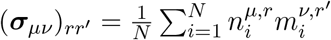 (4) and (5) become:

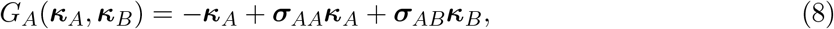

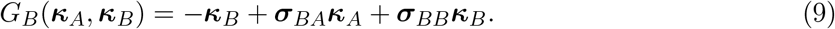

In this case, there are no interactions between tasks if connectivity vectors corresponding to different tasks are mutually orthogonal: ***m***^*µ*^⊥***m***^*ν*^, ***n***^*µ*^⊥***n***^*ν*^, ***m***^*µ*^⊥***n***^*ν*^, for *μ* ≠ *ν*. When the last of these requirements holds, ***σ***_*AB*_ = ***σ***_*BA*_ = 0, so that the dynamics of ***κ***_*A*_ and ***κ***_*B*_ decouple into 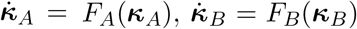. Because activity in directions orthogonal to the task A and B subspaces decays, this satisfies the criteria for simultaneous multi-tasking, with the limitation that single tasks must correspond to linear dynamical systems.

### A.6 Multi-tasking in nonlinear networks: mean-field approach

We now consider nonlinear networks in the large-*N*, mean-field limit. We will again assume that the connectivity vectors for different tasks are orthogonal. This assumption is, in general, stronger than required for topological conjugacy, since some degree of overlap across tasks may distort but not completely reconfigure task-associated flow fields. The amount of overlap that can be tolerated will depend on the specifics of the tasks and the network.

The more significant observation for our present analysis is that, even when the connectivity vectors are orthogonal, the criteria for multi-tasking may not be satisfied in nonlinear networks. In particular, we show that while the dynamics in each task subspace may correctly correspond to those of the desired task, these subspaces may not be attracting.

#### A.6.1 Orthogonal and independent connectivity vectors preserve dynamics within task subspaces

The mean field limit we consider corresponds to assuming that, for each neuron *i*, the components of its connectivity vectors are drawn from a probability distribution, 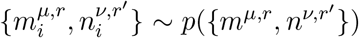), independently across neurons, for *μ, ν* = 1 … *P* and *r, r*^*′*^ = 1 … *R*_*µ/ν*_. Under this assumption, sums of the form 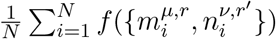) converge to 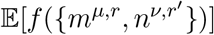 where E is the expectation under the distribution *p*.

Beyond the mean-field assumption, we will further assume that connectivity components are statistically independent across tasks, so that 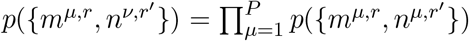, and that each connectivity vector has zero-mean elements, E [*m*^*µ,r*^] = E [*n*^*µ,r*^] = 0. These two assumptions imply that connectivity vectors corresponding to different tasks are orthogonal in the large-*N* limit.

Under these assumptions, we first show that the dynamics restricted to the task A subspace are the same as *F*_*A*_, the dynamics of a single-task network. Let ***M*** ^*µ*^ and ***N*** ^*µ*^ be *N* × *R*_*µ*_ matrices whose *r*th columns are equal to ***m***^*µ,r*^ and ***n***^*µ,r*^, respectively. The dynamics of ***κ***_*A*_ follow

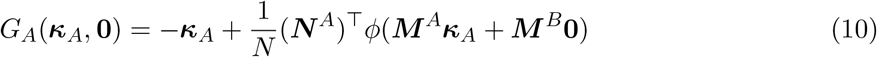

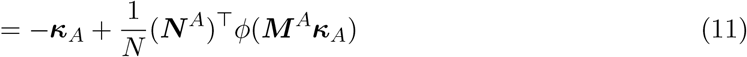

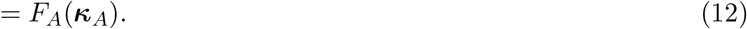

Now we consider the dynamics of ***κ***_*B*_. The fact that the connectivity components for tasks A and B are independent and zero-mean implies that *G*_*B*_ = 0, because

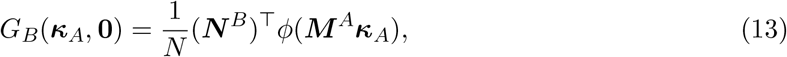

and, for any *r*,

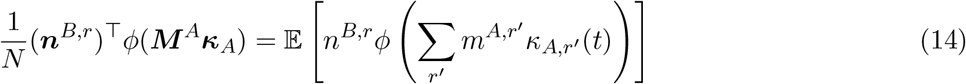

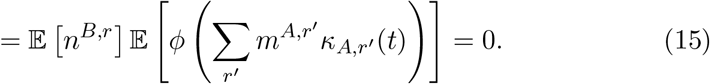

This, along with Eq. (12), implies that, on the task A subspace, 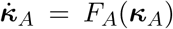and 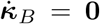. An analogous argument holds for task B and for any finite number of tasks in which all pairs of connectivity components are orthogonal and independent. These dynamics clearly satisfy the topological conjugacy requirement for sequential multi-tasking, with the mapping from single-task to multi-task dynamics being the identity function.

#### A.6.2 Gaussian connectivity vectors lead to transverse instability

The next requirement for multi-tasking is that each task subspace is attracting. We show that, if *p*({*m*^*µ,r*^, *n*^*ν,r*^*′*}), the probability distribution that defines the connectivity vectors, is jointly Gaussian, and tasks involve sustained dynamics, only one subspace satisfies this criterion, so that such networks do not successfully multi-task (as in Fig. 2C). It can be shown that, in this case, the dynamics of ***κ***_*µ*_ follow ^16,21^:

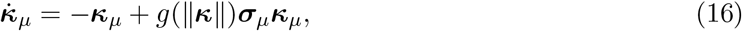

where 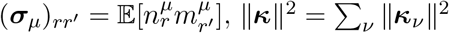 and *g* = E [*ϕ*^*′*^(*x*)] is a scalar function, representing the average neural gain across the network. We assume *ϕ* is nondecreasing, so that *g* ≥ 0. Eq. (16) implies that the transverse stability of task A with respect to task B is determined by the transverse Jacobian (Eq. 7):

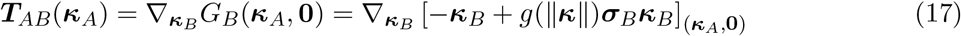

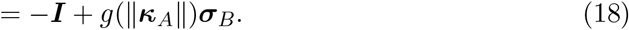

Note that the eigenvectors of Eq. (18) do not depend on ***κ***_*A*_, which implies that local transverse stability determines whether the task A subspace is attracting, as discussed in Sec. A.4. Let *α*(***σ***) denote the largest real part of the eigenvalues of ***σ***. Assuming this quantity is positive, the largest real part of the transverse eigenvalues is then

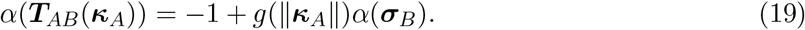

We now argue that, for the tasks we consider, if *α*(***σ***_*B*_) *> α*(***σ***_*A*_), the task A subspace is locally transversely unstable. The argument relies on an assumption that, for every task, *g*(∥***κ***_*µ*_∥)*α*(***σ***_*µ*_) ≥ 1 for at least one value of ***κ***_*µ*_. In Sec. A.6.5, we show that this requirement is necessary for any task with sustained dynamics, i.e., for which ***κ***(*t*) does not decay to 0 for large *t*.

However, since *α*(***σ***_*B*_) *> α*(***σ***_*A*_), for any point at which *g*(∥***κ***_*A*_∥)*α*(***σ***_*A*_) ≥ 1, Eq. (19) is positive, and thus the task A subspace is locally transversely unstable. The same argument implies that if *α*(***σ***_*A*_) > *α*(***σ***_*B*_), the task B subspace is locally transversely unstable. More generally, all task sub-spaces except for those with the largest value of *α*(***σ***_*µ*_) are locally transversely unstable. Avoiding interference due to this instability would require fine-tuning to ensure all *α*(***σ***_*µ*_) are equal. This limitation underlies the task interference shown in Fig. 2.

#### A.6.3 Requirement for transient multi-tasking

This linear analysis can be used to estimate the time interval associated with transient solutions to multi-tasking as in Fig. 3. Suppose an initial condition is set as ***κ***(0) = *c*_*A*_***v***_*A*_ + *c*_*B*_***v***_*B*_. The initial condition has a component along the vector ***v***_*A*_, which lies within the task A subspace, and a small component along ***v***_*B*_ (that is, |*c*_*B*_| ≪ |*c*_*A*_|), the eigenvector in the task B subspace associated with the largest real eigenvalue component as defined above. Suppose that *g*(∥***κ***_*A*_(0)∥)*α*(***σ***_*B*_) > 1.

At small *T*, the magnitude of the projection of ***κ***(*T*) onto the task B subspace is then approximated by |*c*_*B*_| exp((*g*(∥***κ***_*A*_∥)*α*(***σ***_*B*_) − 1)*T*). In this case, for the ratio of the projection onto the task B and task A subspaces to remain below *r* for a time interval of length *T*, the initial condition must satisfy

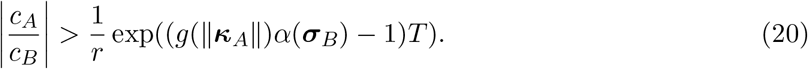

Thus, increasing *T* requires fine-tuning of the initial condition to minimize the component along amplifying directions for task B. The degree of fine-tuning depends exponentially on *g*(∥***κ***_*A*_∥)*α*(***σ***_*B*_)−1. This analysis is formally valid in the limit of small *r* and |*c*_*B*_|.

#### A.6.4 Requirement for multi-tasking through effective connectivity modulation

Eq. (19) can also be used to determine the required adjustment of the connectivity strength *a*_*A*_ (main-text Eq. 3) to assure transverse stability of the task A manifold when *α*(***σ***_*B*_) > α(***σ***_*A*_):

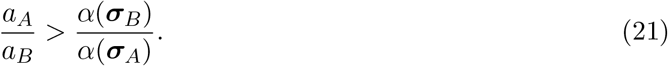

More generally, for networks that perform more than two tasks, the largest required adjustment equals (max_*µ*_ *α*(***σ***_*µ*_))/(min_*µ*_ *α*(***σ***_*µ*_)).

#### A.6.5 Condition for sustained activity in networks with Gaussian connectivity

For a network with Gaussian-distributed connectivity vectors, ***κ*** follows ^16,21^:

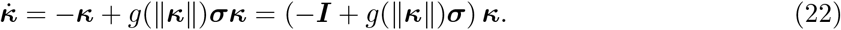

We assume *g* ≥ 0. We will show that if *g*_max_ *α*(***σ***) < 1, where *g*_max_ = max_***κ***_ *g*(∥***κ***∥) and *α*(***σ***) is the largest real part of the eigenvalues of ***σ***, then ***κ*** = **0** is globally attracting.

Because the nonlinearity enters only as a scalar multiplying ***σ***, the time-dependent matrix (−***I*** + *g*(∥***κ***(*t*)∥)***σ***) multiplying ***κ***(*t*) in Eq. (22) commutes for any two timepoints, and the solution can be written

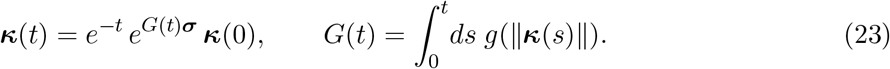

Note that 0 ≤ *G*(*t*) ≤ *g*_max_ *t*. This implies that ∥*e*^*G*(*t*)***σ***^∥ grows no faster than *e*^*βt*^ for large *t*, for any *β* ∈ (*g*_max_ *α*(***σ***), 1). Thus, for some constant *C*,

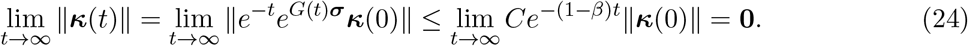

#### A.6.6 Conjugate dynamics across thresholds

For a network with Gaussian-distributed connectivity vectors and a rectified-linear activation function with threshold *θ*_*µ*_, that is, *ϕ*(*x*) = max(*x* − *θ*_*µ*_, 0), ***κ***_*µ*_ follows:

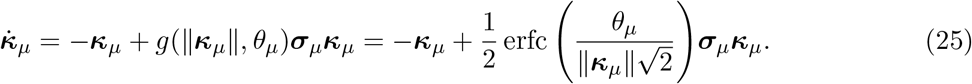

The second equality is derived from 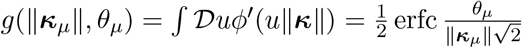.

If ***σ***_*µ*_ = ***σ***_*ν*_, the dynamics of ***κ***_*µ*_ are conjugate to ***κ***_*ν*_ as long as *θ*_*µ*_ and *θ*_*ν*_ have the same sign, which is verified by the homeomorphism ***κ***_*ν*_ = *θ*_*ν*_***κ***_*µ*_/*θ*_*µ*_ that satisfies the conjugacy condition. This simple equivalence arises because of the threshold appearing only as a scale parameter in Eq. (25), so that flow fields associated with ***κ***_*µ*_ and ***κ***_*ν*_ are identical up to scaling by *θ*_*ν*_/*θ*_*µ*_. Thus, somewhat surprisingly, the threshold does not affect the qualitative dynamics of Gaussian networks with fixed ***σ*** unless a sign change occurs.

### A.7 Effects of non-Gaussian connectivity on multi-tasking

In this section, we show that non-Gaussianity in the distributions of connectivity vector elements can change the qualitative behavior of multi-tasking solutions. To simplify the derivation, we assume each task is associated with one-dimensional bistable dynamics, like those of Task B in Fig. 3, and that *g*_*I*_ = 0. In this case, the synaptic weight matrix is:

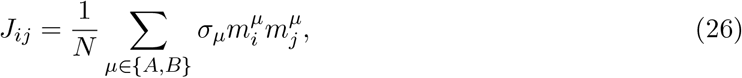

where 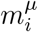 is a random i.i.d. draw from the probability distribution *P*(*m*^*µ*^). We assume the probability distribution is the same for all tasks so that *P*(*m*^*µ*^) = *P*(*m*) for all *μ*, symmetric about the origin, and normalized to have unit variance (Fig. S3A). The dynamics of this network’s task variables are then:

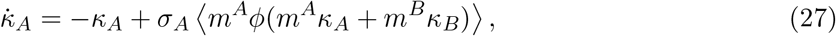

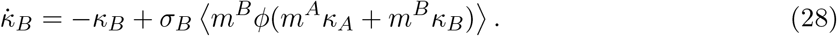

The activation function is threshold-linear, *ϕ*(*x*) = max(*x* − *θ, 0*), with *θ* < 0 so that activity is superthreshold. The network has a trivial fixed point at which *κ*_*A*_ = *κ*_*B*_ = 0. Amplification with *σ*_*µ*_ > 1 is required to generate nontrivial dynamics, leading to fixed points where 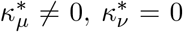 for *μ* ≠ *ν*. We further require *σ*_*µ*_ < 2, which ensures bounded dynamics. We focus on a nontrivial fixed point along the *κ*_*A*_ subspace, which is obtained by setting 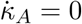:

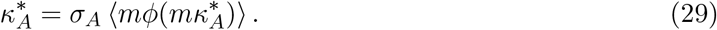

We wish to assess how the transverse stability of this fixed point depends on the distribution *P*(*m*) and on the ratio of *σ*_*A*_ and *σ*_*B*_. The stability is determined by the eigenvalue

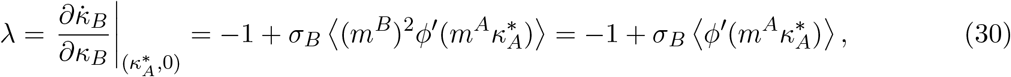

where the second equality comes from the independence of *m*^*B*^ and *m*^*A*^ and the unit variance assumption.

**Figure S3:**
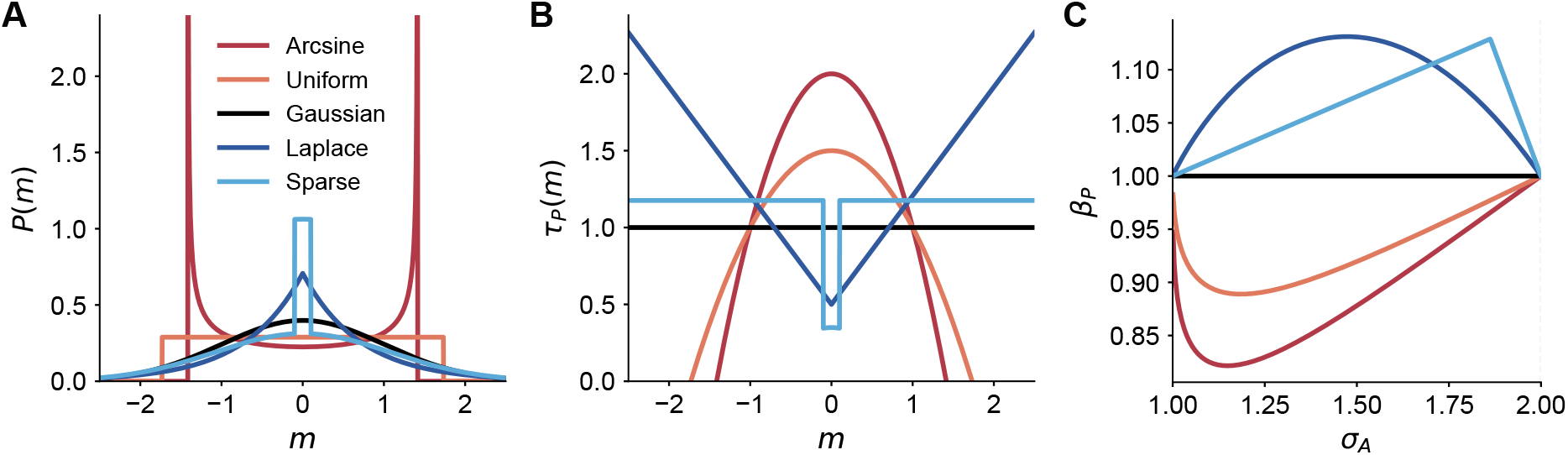
Effect of connectivity pattern distribution on stability. **A** Probability density function *p*(*m*) for five different symmetric distributions with unit variance. The arcsine distribution arises when the connectivity pattern is given by the cosine of a uniformly distributed angle: 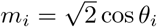 for random uniform *θ*_*i*_. The “sparse” distribution is a mixture of a uniform distribution |*m*_*i*_| < 0.1 with weight 0.15, and a Gaussian distribution normalized such that the total variance is one. **B** Stein kernel for the different distributions. **C** Factor *β*_*P*_ for the different distributions as *σ*_*A*_ is increased. Threshold *θ* = −1.

To express Eq. (30) in terms of the ratio of *σ*_*A*_ and *σ*_*B*_, we derive an expression for *σ*_*A*_ based on the fixed point equation (Eq. 29) using Stein’s identity:

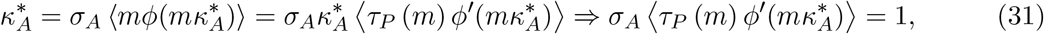

where we have defined the Stein kernel of the distribution 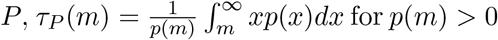 (Fig. S3B). We then define the factor

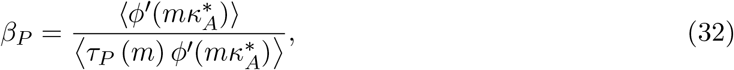

which is determined by the distribution *P*(*m*) and *σ*_*A*_. The transverse eigenvalue can then be expressed in terms of this factor:

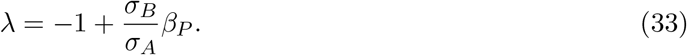

The Stein kernel for a Gaussian distribution is constant, *τ*_*N*_ (*m*) = 1, implying *β*_*P*_ = 1. This is commonly known as Stein’s lemma and often used in analyses of low-rank recurrent neural networks with Gaussian weights^16,21,30^. The argument of Sec. A.6.2, that the task subspace corresponding to the smaller value of *σ*_*µ*_ is transversely unstable, is an example of this.

Non-Gaussian distributions that are unimodal and concentrated around 0, such as the Laplace distribution, have Stein kernels *τ*_*P*_ (*m*) that increase with |*m*|, implying that *β*_*P*_ > 1 when 1 < *σ*_*A*_ < 2 (Fig. S3C). In contrast, distributions less concentrated around 0, such as the uniform distribution or the arcsine distribution that arises from cosine-distributed weights, have Stein kernels that decrease with |*m*|, implying *β*_*P*_ < 1 when 1 < *σ*_*A*_ < 2.

Suppose *σ*_*A*_ < *σ*_*B*_. If *P*(*m*) were Gaussian, the task A subspace would be transversely unstable. However, for distributions with *β*_*P*_ < 1, *λ* may be negative so that task A is stable even in this case. The required deviation from Gaussianity is determined by the ratio 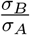, since the stability condition requires 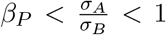. Thus, distributions that are more dispersed than the Gaussian, such as uniform or arcsine distributions, can exhibit multistability and support sequential multi-tasking (e.g. Fig. 4).

On the other hand suppose *σ*_*A*_ > *σ*_*B*_, which would imply transverse stability of the task A subspace if *P*(*m*) were Gaussian. Non-Gaussian distributions that concentrate around the origin, such that 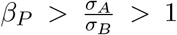, can destabilize both the task A and task B subspaces in this case. We found that when this occurs, the bounded dynamics converge to an attracting simultaneous multi-tasking manifold with nonzero projections onto both *κ*_*A*_ and *κ*_*B*_.

In total, these results show that non-Gaussianity of *P*(*m*) can either increase or decrease transverse eigenvalues, depending on the distribution’s degree of dispersion. While the derivation was performed for one-dimensional task subspaces, we expect the qualitative results to hold for higher dimensions. The specific result that connectivity patterns whose distribution concentrates around zero generate simultaneous multi-tasking solutions motivated us to introduce connectivity barcodes, defined in the next section, as a tractable way to build multi-tasking networks.

### A.8 Connectivity barcodes

In this section, we develop a mean-field theory for networks with connectivity barcodes that describes when rule inputs can select tasks sequentially or simultaneously. The previous section showed that when the distributions of the connectivity vector elements are sparse, single-task solutions tend to be destabilized. As described in the main text, one way to instantiate this mechanism is to attach a sparse barcode to each connectivity component, which means that the connectivity vectors are no longer jointly Gaussian. Let 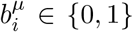 be independent Bernoulli variables with mean *f*, and define

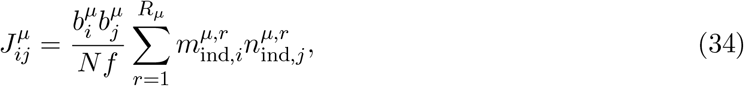

where 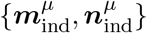 are jointly Gaussian as in Sec. A.6.2. The factor 1/*f* ensures that the single-task dynamics are invariant across choices of *f*. The low-dimensional variables of each task are defined by the projection

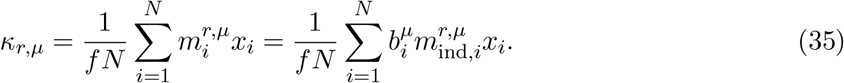

The single-task dynamics that would be produced if 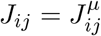 are given by Eq. (16):

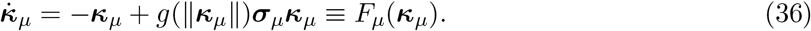

The mean-field dynamics for two tasks now each include a linear combination of two nonlinear terms:

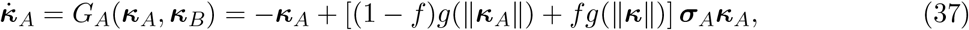

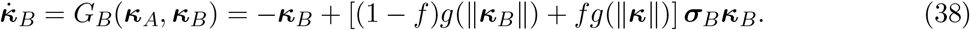

The nonlinear function *g* is defined by the Gaussian integral *g*(∥***y***∥) = Du*ϕ*^*′*^ (*u*∥***y***∥), where ∥·∥ denotes the Euclidean norm, and corresponds to the average gain of neurons within the population representing the variables ***y***. The first term in each bracket comes from neurons for which 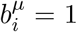 for only one task, and the second comes from neurons for which 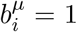 for both tasks.

First, observe that *G*_*A*_(***κ***_*A*_, **0**) = *F*_*A*_(***κ***_*A*_) and *G*_*B*_(**0, *κ***_*B*_) = *F*_*B*_(***κ***_*B*_), meaning that dynamics within each task subspace are conjugate to the desired task variable dynamics. Thus, the topological conjugacy requirement for multi-tasking is satisfied.

As *f* → 0, Eqs. (37) and (38) decouple, and thus the network simultaneously multi-tasks. For the examples we consider, this transition occurs when *f* is small enough that all single-task subspaces lose transverse stability and lead to simultaneous multi-tasking. We first calculate this transition point, and then show how rule inputs can select individual tasks to enable sequential multi-tasking.

#### A.8.1 Barcodes produce single-task transverse instability and enable simultaneous multi-tasking

We will show that single-task subspaces are transversely unstable for all tasks when *f* is small enough. For two tasks A and B, the transverse Jacobian associated with task B’s dynamics on the task A subspace is

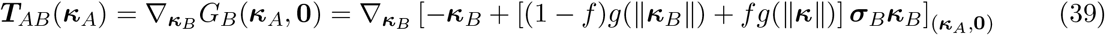

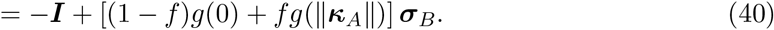

The maximum real part of the Jacobian’s eigenvalues is therefore

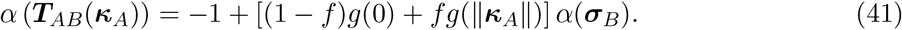

We assume that *g*(0)*α*(***σ***_*B*_) > 1, which ensures absence of a stable trivial fixed point at the origin. Then, for any point ***κ*** = (***κ***_*A*_, **0**) on the task A subspace for which *g*(∥***κ***_*A*_∥) < *g*(0), there is a value of *f* below which this point is transversely unstable. The assumption *g*(∥***κ***_*A*_∥) < *g*(0) is true for networks in which the activation function threshold *θ* < 0, which is necessary for sustained activity. This value of *f* is given by

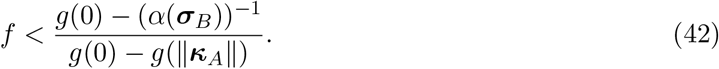

We now consider the specific case in which each task subspace has an attractor 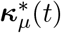 on which 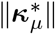 is constant (for example, a fixed point or a circular limit cycle), and we evaluate transverse stability on this attractor. In this case, 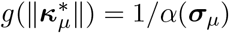. This yields the criterion

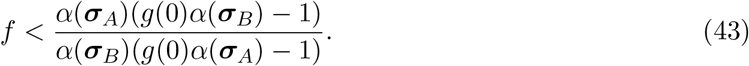

An identical argument holds for the task B subspace. In simulations, we have found that when all single task solutions are transversely unstable and the dynamics do not grow unboundedly (e.g., with a bounded nonlinearity, or *α*(***σ***) < 2 for threshold-linear neurons) there is a manifold that supports the simultaneous performance of tasks A and B. For networks with more than two tasks, if Eq. 42 holds for all pairs of tasks, all task subspaces are transversely unstable.

#### A.8.2 Rule inputs in barcoded networks enable sequential multi-tasking

We have shown that barcodes produce simultaneous multi-tasking in networks without external input. We now show that it is possible to construct networks with barcodes in which rule inputs select specific tasks, enabling either sequential or simultaneous multi-tasking. We assume that the activation function is rectified-linear, *ϕ*(*x*) = max(*x* − *θ*, 0), but, unlike the previous section, we assume that *θ* > 0, so that in the absence of an input, the network is silent. We then apply a rule input proportional to the barcode of task A:

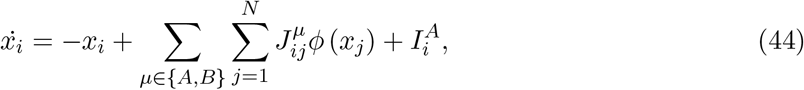

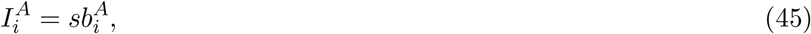

**Figure S4:**
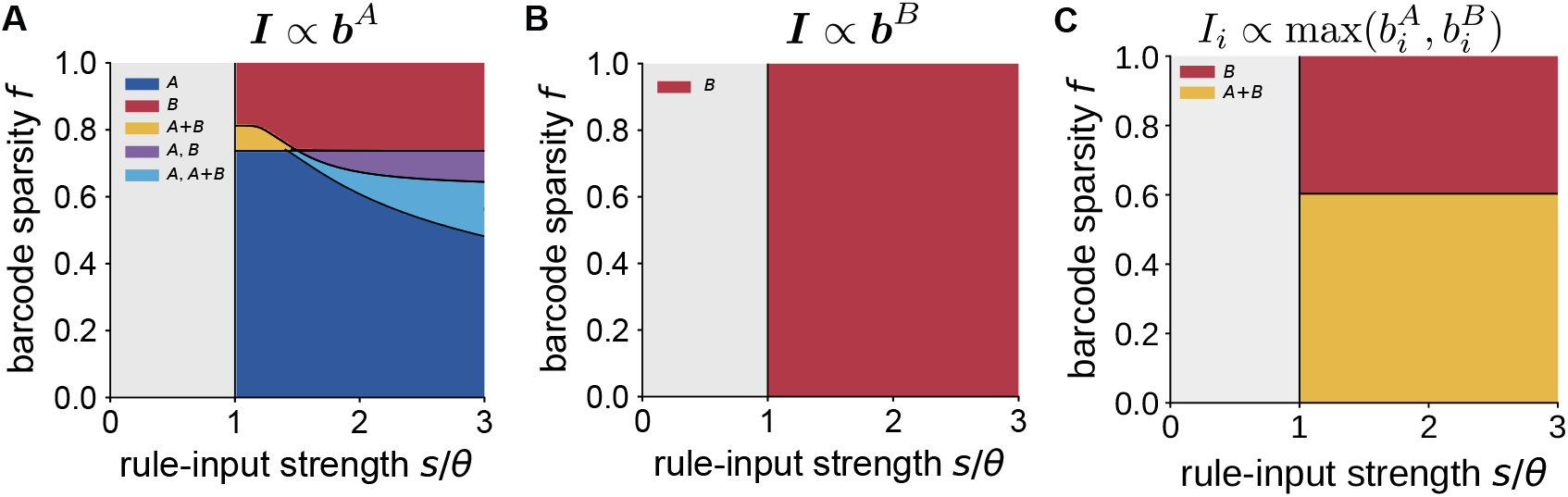
Barcoded rule input in a network with *α*(***σ***_*B*_) *> α*(***σ***_*A*_). **A** Phase diagram showing stable task manifolds as a function of the barcode sparsity and the strength of the rule input. The input is proportional to the task A barcode. If Eq. 51 is satisfied, task A is selected (blue region). Strong inputs may lead to additional stable manifolds. A+B denotes simultaneous multi-tasking. A, B denotes multistability of task A and B solutions. A, A+B denotes multistability of a task A solution and a simultaneous multi-tasking solution. **B** Similar to panel A, when the input is proportional to the task B barcode. **C** Similar to panels A and B, when the input is given by both task’s barcodes. Task parameters: *α*(***σ***_*A*_) = 1.4 and *α*(***σ***_*B*_) = 1.9.

Activity ***x***(*t*) can be parameterized with task variables ***κ***_*µ*_(*t*), which represent projections along the vectors {***m***^*µ,r*^} as in Eq. 3, together with an additional variable *κ*_*I*_ to parameterize activity projected onto the vector ***b***^*A*^. The dynamics of these latent variables are

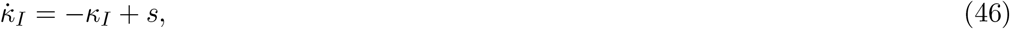

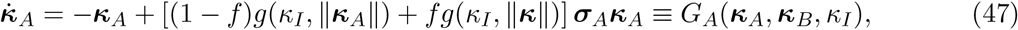

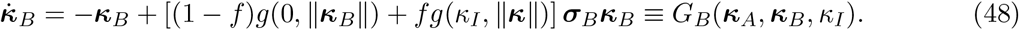

The nonlinear function *g* is defined by the Gaussian integral *g*(*μ*, ∥***y***∥) = *Duϕ*^*′*^ (*μ* + *u*∥***y***∥), where *μ* describes the average current to each neural population.

##### Activation of a selected task

We first show that, as long as the input strength *s* is larger than the single neuron threshold, *s* > *θ*, and the barcode is sparse enough, the multi-tasking network generates an attracting single-task subspace. We assume that the dynamics *F*_*µ*_ associated with the performance of task *μ* are again given by Eq. (36), corresponding to the dynamics of a single-task network. The activation function for neurons in this single-task network has a threshold that we denote by *θ*_1_ < 0.

In the barcoded network, the dynamics on the task A subspace are determined by Eq. (47) when ***κ*** = (***κ***_*A*_, **0**). In this subspace, the dynamics do not depend on the value of *f* and have the same form as the desired single-task dynamics *F*_*A*_, except that the effective activation function threshold is different and given by *θ* − *s*. The result of Sec. A.6.6 implies that as long as *s* > *θ*, the barcoded network’s dynamics are conjugate to those defined by *F*_*A*_. Thus, a relatively unrestrictive condition on the magnitude of the rule input is sufficient to produce the desired dynamics.

Next, to assess stability of the task A subspace, we evaluate the largest real part of the transverse eigenvalues:

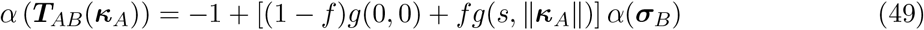

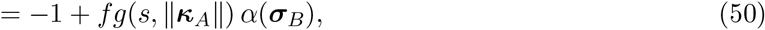

where the second line comes from the assumption of a positive threshold of the activation function, implying *g*(0, 0) = 0. Because *g* ≤ 1, transverse stability is therefore always guaranteed if *f* < 1/*α*(***σ***_*B*_). More generally, in networks that perform more than two tasks, if *f* < 1/*α*(***σ***_*µ*_) for all *μ*, all task subspaces are transversely stable. Thus, we have established topological conjugacy and attractiveness of task subspaces when a rule input is provided, if *f* is small enough.

We now derive a less restrictive bound on *f* for specific single-task dynamics, such as fixed points or limit cycles, that are generated by an instability when ***κ***_*µ*_ approaches zero. For such tasks, the dynamics stabilize when the amplifying dynamics are balanced by the average neuronal gain: *g*(∥***κ***_*µ*_∥)*α* (***σ***_*µ*_) = 1. Substituting this into Eq. (50) leads to the following condition on *f*:

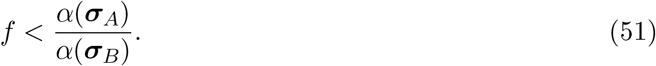

An example phase diagram for a network with *α*(***σ***_*A*_) *< α*(***σ***_*B*_) and receiving a rule input proportional to the task A barcode is shown in Fig. S4A. The condition of Eq. (51) is satisfied in the blue, cyan, and purple regions. In these regions an attracting task A subspace exists (although in the purple and cyan regions, another multistable solution also exists, see next section). On the other hand, the same network receiving a task B barcode rule input only performs task B, since *α*(***σ***_*A*_) < α(***σ***_*B*_) (Fig. S4B).

##### Complete phase diagram

We next analyze under what situations a network receiving a rule input proportional to the task A barcode also generates dynamics in the task B subspace (Fig. S4A, red, yellow, purple, and cyan regions). These solutions can be viewed as spurious states in which an input intended to select task A can select another task or combination of tasks. As in the derivation of Eq. (51), we assume that each task is associated with an attractor 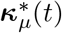 on which 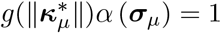. Such solutions can be characterized analytically in the limit *s* = *θ* + *ϵ, ϵ* → *0*.

Based on Eq. (48), when *s* is above threshold but small enough such that 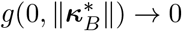, activity in the task B subspace is nondecaying only if *f* > 1*/α*(***σ***_*B*_). Transverse stability of the task B subspace is determined by:

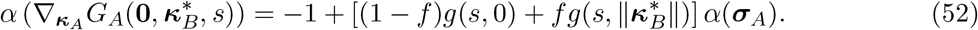

The gain factor *g*(*s*, 0) = 1 for *s* > *θ*. Additionally, the task dynamics impose 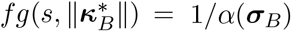, so that task B is stable only if

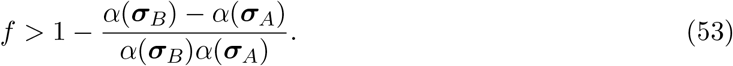

Note that this can only be satisfied if *α*(***σ***_*B*_) *> α*(***σ***_*A*_). This shows that, if task B is more amplifying than task A and *f* is too large, a rule input intended to select task A can instead select task B (Fig. S4A, red region).

Next, we examine whether a simultaneous multi-tasking solution exists, i.e. 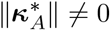 and 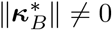. Using Eq. (48), such a solution requires *fg*(*s*, ∥***κ***^*\**^∥) = *α*(***σ***_*B*_)^−1^. Introducing this result into Eq. (47), we obtain the condition (1 − *f*)*g*(*s*, ∥***κ***_*A*_∥) = *α*(***σ***_*A*_)^−1^ − *α*(***σ***_*B*_)^−1^. Such solutions require 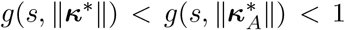. Combining these three expressions, we obtain a range of sparsity values for which the simultaneous multi-tasking solution exists (Fig. S4A, yellow region):

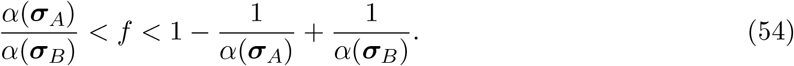

This range coincides with the range in which both the task A and B subspaces are unstable (Eqs. 51 and 53). Evaluation of the Jacobian associated with this simultaneous multi-tasking solution shows that, if the solution exists, it is always stable.

The full phase diagram, for arbitrary *s*, can be assessed numerically. As the input strength increases further from *s* ≳ *θ*, multistable phases also appear, where both the task A and task B attractors are stable, or even multistable phases where both a single task A attractor and a simultaneous task A and B attractor are stable (Fig. S4A, purple and cyan regions). These phase boundaries are calculated by numerically finding roots to the existence and transverse stability equations.

##### Combination of rule inputs for simultaneous multi-tasking

We have shown it is possible to select task A (Fig. S4A, blue region) or task B (Fig. S4B, red region) with a rule input proportional to the task barcodes. To select tasks *μ* and *ν* simultaneously, we provide a rule input that combines two barcodes, 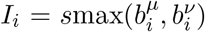. If the network performs only two tasks A and B, the input is received by all neurons with 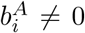 or 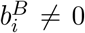 homogeneously, corresponding to an effective change in these neurons’ mean input. Therefore, the network simultaneously multi-tasks if *f* is below the critical value given by Eq. (42) (Fig. S4C).

#### A.8.3 Barcodes with non-binary elements

We now consider a barcode model with

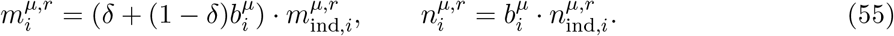

If *δ* > *0*, neurons that do not drive the dynamics of a task (that is, neurons *i* for which 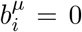 and thus 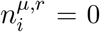) are nonetheless tuned to that task. The magnitude of *δ* controls the strength of this tuning, with *δ* = 0 reducing to the binary barcode model of the previous section and *δ* = 1 corresponding to equal tuning to all tasks. The low-dimensional variables are rescaled to account for this parameter so that the single-task dynamics remain unchanged:

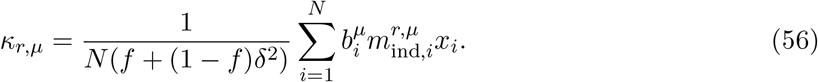

The dynamics in a two-task network, analogous to Eqs. (37) and (38), are:

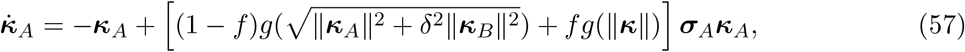

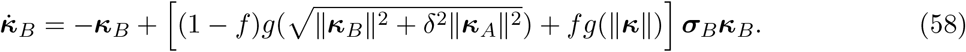

We consider a network with *θ* < 0, and 1 < *α*(***σ***_***B***_) < *α*(***σ***_***A***_) < 2. The maximum real part of the transverse eigenvalues for the task A subspace is

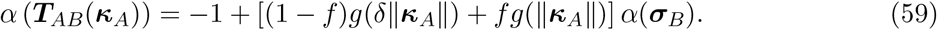

For tasks with constant norm, 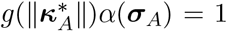. Under this assumption, the sparsity criterion that ensures the single-task solution is unstable is:

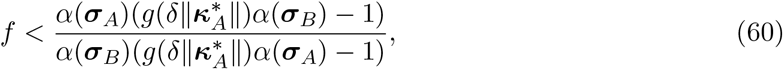

which can be solved numerically. As in previous sections, when the task A subspace is transversely unstable, a simultaneous multi-tasking solution emerges.

The criterion in Eq. (60) shows that there is a trade-off between the selectivity of neurons to tasks whose dynamics they do not drive, described by *δ*, and the barcode sparsity *f*. The maximum *f* at which simultaneous multi-tasking is achieved corresponds to *δ* = 0, and is determined by Eq. (43). As *δ* increases, *f* must decrease to permit simultaneous multi-tasking. In the limit *f* → 0, the maximum *δ* that permits simultaneous multi-tasking is:

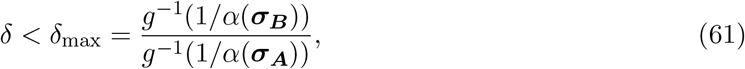

where *g*^−1^ denotes the inverse of the average gain function.

### A.9 Global inhibition in threshold-linear networks

Unbounded single neuron activation functions, such as rectified-linear, can lead to instability for strongly amplifying dynamics. In particular, if *α* (***σ***_*µ*_) > 2, amplification cannot be compensated by the gain term *g*(∥***κ***_*µ*_∥), which in this case has a lower bound of 1/2 for *θ* < 0. One solution to such instability is the addition of global inhibition.

Following the main text, we consider an inhibitory population that integrates activity across all neurons in the network and provides negative feedback with strength *g*_*I*_ (Eq. 2). The dynamics of the latent variables, parameterized in Eq. (3), are then:

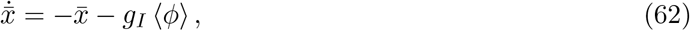

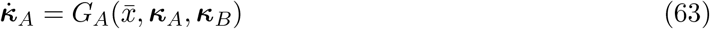

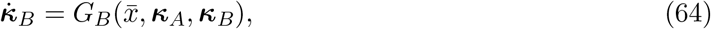

where ⟨*ϕ*⟩ indicates the average firing rate.

For networks with inhibition, the trivial fixed point is defined by ***κ***_*A*_ = ***κ***_*B*_ = **0** and 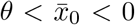, where 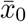 satisfies 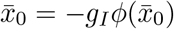. At this fixed point, the variable 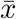 changes the effective threshold of the nonlinearity from 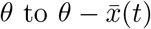. Nevertheless, since 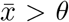, the effective threshold does not change sign, and the result of Sec. A.6.6 implies that the dynamics in the vicinity of the trivial fixed point remain conjugate to the dynamics without inhibition. A similar effect is observed for task-related dynamics, as long as 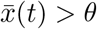. For the fixed-point and limit-cycle tasks we have considered, 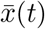 is larger than the threshold if *α* (***σ***_*µ*_) < 2, so that multi-tasking solutions are qualitatively unchanged by the presence of inhibition.

However, if 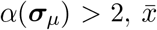 becomes small enough that the effective threshold changes sign to compensate the amplifying task dynamics. In this parameter range, task dynamics are stabilized by global inhibition, and this affects solutions qualitatively. For example, for non-Gaussian connectivity vectors, the factor *β*_*P*_ (Eq. 32; Fig. S3) crosses the unity line when *α*(***σ***_*µ*_) = 2. Therefore, a non-Gaussian distribution that generates multistable task manifolds without inhibition will generate a simultaneous multi-tasking manifold in the inhibition-dominated regime. Conversely, for non-Gaussian distributions that lead to simultaneous multi-tasking, such as connectivity barcodes, the inhibition-dominated regime leads instead to multistability.

For barcoded connectivity, intuition is provided by noting that in the absence of inhibition, barcodes lead to simultaneous multi-tasking when they are sparse enough because specialized neural populations encode different tasks with little interference. The intuition is reversed in networks stabilized by global inhibition: whenever one population is active, feedback inhibition silences all others, leading to multistability.

In conclusion, global inhibition can stabilize networks with threshold-linear activation functions and affects their multi-tasking properties. In particular, changes in inhibition can cause a network that simultaneously multi-tasked to sequentially multi-task via multistable dynamics. Alternative inhibitory mechanisms, such as task-specific inhibition, will also likely have different effects on multi-tasking.

### A.10 Effects of activation function

Most theories of nonlinear networks with low-rank connectivity have used neuron models with bounded sigmoid activation functions such as *ϕ*(*x*) = tanh(*x*), though many results do not depend on this choice. Here, we have focused on threshold-linear functions, which are consistent with the presence of a spike threshold and inter-neuronal communication via nonnegative firing rates.

The mean-field description of networks with low-rank connectivity applies to any pointwise non-linear activation function. In this description, the dynamics of single-task networks are given by Eq. (16). The scalar nonlinear function *g*, which represents the average gain across the network, depends on the nonlinearity. However, for any non-decreasing activation function, including threshold-linear and sigmoid functions, Gaussian connectivity vectors lead to transverse instability.

Our results on the effects of non-Gaussian connectivity in Sec. A.7 also extend to networks with sigmoid activation functions. Indeed, whether the factor *β*_*P*_ (Eq. 32) is above or below one, which determines whether networks exhibit multistability or support simultaneous multi-tasking, depends on how *ϕ*^*′*^(*mκ*^*\**^) behaves as |*m*| increases. For both hyperbolic tangent and threshold-linear activation functions in networks with 1 *< α*(***σ***) < 2, the gain decreases as |*m*| increases. Nevertheless, as *α*(***σ***) approaches 2, for the former and unlike the latter case, the factor *β*_*P*_ does not approach one, but remains above or below one depending on the distribution.

The sparsity criterion that determines when networks simultaneously multi-task (Eq. 43) also holds for networks with sigmoid activation functions. An important difference between networks with different activation functions appears when using rule inputs in barcoded networks (Sec. A.8.2). For threshold-linear networks, rule inputs drive specific neuronal subpopulations above threshold, while other neurons remain below threshold. Networks with sigmoid activation functions do not have a threshold, but can implement a similar mechanism. Strong inputs, both positive and negative, decrease the gain of a population by driving its neurons towards saturation.

One implementation of rule inputs in networks with sigmoid activation functions provides a background input negatively correlated with all barcodes: ***I***_0_ = −*s*_0_ ∑_*µ*_ ***b***_*µ*_, which broadly suppresses task-related dynamics. To select task *μ*, an additional rule input correlated with task *μ*’s barcode is provided, ***I***_*µ*_ = *s****b***_*µ*_. The complete quantitative phase diagram, and its behavior as *s* varies, will depend on the specific activation function.

## Footnotes

† In terms of *G* and *F*_*µ*_, for example, 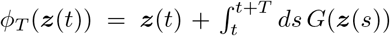 and 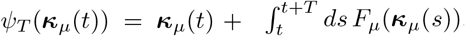.

† Counterexamples can be constructed by considering the system 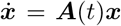, and interpreting *t* as the location along a one-dimensional manifold and ***x*** as the transverse dynamics. Josić & Rosenbaum ^75^ provide a didactic account of several counterintuitive examples in which ***A***(*t*) has eigenvalues with negative real part for all *t*, yet ***x*** = **0** is unstable. The examples rely on the eigenvectors of ***A***(*t*) changing with *t*.

